# NTpred: A robust and precise machine learning framework for in-silico identification of Tyrosine nitration sites in protein sequences

**DOI:** 10.1101/2023.02.18.529069

**Authors:** Sourajyoti Datta, Muhammad Nabeel Asim, Andreas Dengel, Sheraz Ahmed

## Abstract

Post-translational modifications (PTMs) either enhance a protein’s activity in various sub-cellular processes, or degrade their activity which leads towards failure of intracellular processes. Tyrosine nitration (NT) modification degrades protein’s activity that initiate and propagate various diseases including Neurodegenerative, Cardiovascular, Autoimmune diseases, and Carcinogenesis. Identification of NT modification support development of novel therapies and drug discoveries for associated diseases. Identification of NT modification in biochemical labs is expensive, time consuming, and error-prone. To supplement this process, several computational approaches have been proposed. However these approaches remain fail to precisely identify NT modification, due to the extraction of irrelevant, redundant and less discriminative features from protein sequences. The paper in hand presents NTpred framework competent in extracting comprehensive features from raw protein sequences using four different sequence encoders. To reap the benefits of different encoders, it generates four additional feature spaces by fusing different combinations of individual encodings. Furthermore, it eradicates irrelevant and redundant features from eight different feature spaces through a Recursive Feature Elimination process. Selected features of four individual encodings and four feature fusion vectors are used to train eight different Gradient Boosted Tree classifiers. The probability scores from the trained classifiers are utilized to generate a new probabilistic feature space, that is utilized to train a Logistic Regression classifier. On BD1 benchmark dataset, the proposed framework outperform existing best performing predictor in 5-fold cross validation and independent test evaluation with combined improvement of 13.7% in MCC and 20.1% in AUC. Similarly, on BD2 benchmark dataset, the proposed framework outperform existing best performing predictor with combined improvement of 5.3% in MCC and 1.0% in AUC.

## Introduction

Proteins are essential macro-molecules for human beings and other living organisms. They perform various intracellular activities, such as metabolic reaction catalyzation, DNA replication, stimuli response, and physical processes, such as, providing structure to cells and organisms, and act as transport molecules within various biological systems [8]. Proteins undergo diverse types of Post-translational Modifications (PTMs), such as, Phosphorylation, Acetylation, Ubiquitylation, and many more, that alter the structure of proteins [9]. These modifications improve the activity of proteins in various cellular processes such as, regulation of genetic expression, activation of genes, DNA repair and cell cycle progression, and different physical processes such as signal transduction, chromatin stability, protein–protein interaction and nuclear transport [9]. Contrarily, some modifications such as Nitration, S-nitrosylation and S-palmitoylation adversely affect the activity of proteins in various aforementioned processes [10].

To date, more than 620 different types of PTMs have been identified, that influence the biology of the cell by affecting the functional diversity of the proteome [10]. Among these modifications, Tyrosine nitration modification is considered one of the most important due to its significant involvement in the initiation and propagation of numerous diseases [11–14]. Tyrosine nitration is a PTM where a covalent bond between the Tyrosine and a nitro group (*NO*_2_) produces Nitrotyrosine (NT) [11–14]. Figure 1 graphically illustrates the two stage process of Tyrosine nitration modification, where, in the first stage, Tyrosine undergoes oxidation to form the Tyrosyl Radical, and in the second stage, the radical undergoes nitration to form 3-nitrotyrosine.

**Fig. 1.**
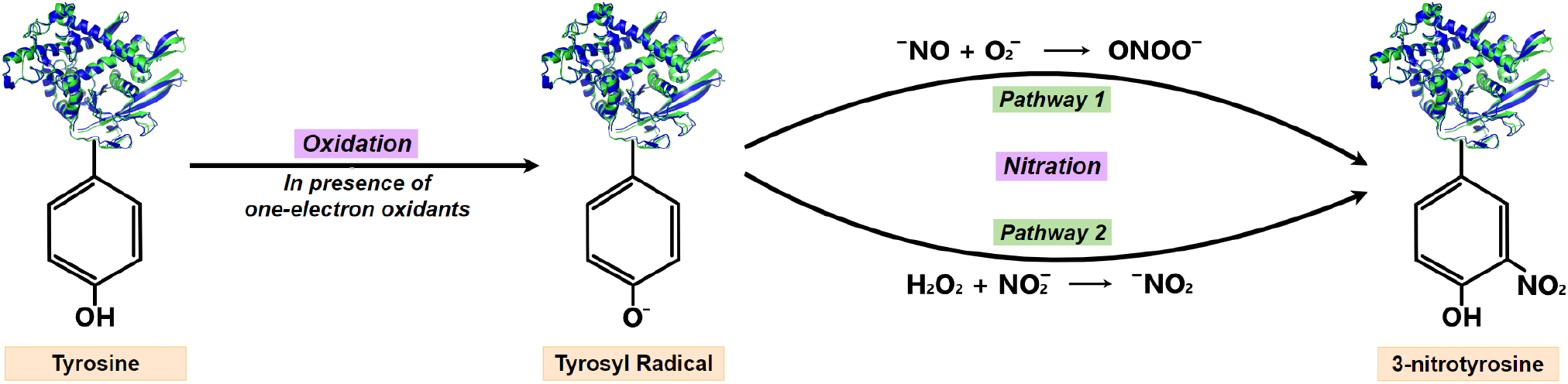
Biochemistry of tyrosine nitration in proteins. It occurs in two stages. First, the Tyrosine undergoes oxidation in the presence of one-electron oxidants, such as ^−^*NO_2_, ^−^OH* or 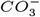, that produces the Tyrosyl Radical. Second, addition of Nitrite (^−^*NO*_2_) to the Tyrosyl Radical by a nitrating agent (*OONO^−^* or ^−^*NO*_2_) produces 3-nitrotyrosine.

Tyrosine nitration based structural alteration of proteins negatively impacts the activity of proteins in various processes, such as signal transduction in cells, energy production in mitochondria, antioxidant defense, and stimulation of the immune system, and sometimes renders a protein completely inactive [11–14]. It inhibits the signal transduction pathways in cells, thus impeding cellular responses [12]. The causal nitration agents have been identified in various disease conditions, such as, Nitric Oxide (NO) produced at a high rate in inflammatory conditions, Nitrite ion (NO2-) greatly increased in systemic inflammatory disorders (sepsis, gastroenteritis, & hemolytic diseases), and abnormal elevation of copper ion (Cu2+) and free heme catalysts in type 2 diabetes mellitus, neurological disorders, and severe hemolytic diseases [12, 14]. Furthermore, NT has been identified in large number of pathological conditions, such as, Neurodegenerative diseases (Parkinson’s and Alzheimer’s, degeneration of dopamine neurons, cerebral ischemia and edema), Cardiovascular diseases, Autoimmune diseases (Rheumatoid Arthritis, Systemic Lupus Erythematosus) and in Carcinogenesis (Breast, Esophageal and Gastric cancer; Colorectal, Squamous cell, Adeno- and Cholangial carcinoma) [12, 14]. Tyrosine nitration is a selective process since it depends upon the accessibility of the Tyrosine residues to the nitrating agents, for example, Tyrosine residues exposed on the surface of proteins can become targets and most nitrated Tyrosine s are in the vicinity of a site which generates nitrating agents [11–14]. Therefore, identification of protein nitration can be employed to recognize onset and progression of the associated diseases, and act as surrogate markers for the design of novel clinical interventions, such as therapeutic strategies and drugs [13, 14].

Traditionally, Tyrosine nitration modification of proteins is detected in biochemical labs through diverse types of in-vivo, ex-vivo, and in-vitro methods, such as, mass spectrometry, histochemical analysis and chromatography [5, 6]. However, due to the sparse occurrence of endogenously nitrated sites in proteins, detections are inefficient [5]. Although incorporation of prior immunoprecipitation techniques in spectrometry improves the detection of Tyrosine nitration sites, however ensembling of both approaches makes the detection process even more complicated [5]. Hence, wet lab experimental approaches tend to be technically challenging, labor intensive, time consuming, expensive and have biases in proteome wide identification [1–7]. Public availability of annotated Tyrosine nitration modification sites in different databases, such as PhosphoSitePlus, ProteomeScout, Human Protein Reference Database, PROSITE, Protein Information Resource (PIR), dbPTM, SysPTM2.0, Uniprot and O-GlcNAc Database [62–70], enables the development of computational approaches that take advantage of Machine Learning and Deep Learning algorithms. These computational approaches can be easily developed, are comparatively much less labor intensive, economical and reusable.

Following the success of machine learning and deep learning approaches in various application areas of bioinformatics, to date researchers have developed 7 predictors with the capability to identify tyrosine nitration sites in protein sequences [1–7]. The working paradigm of these predictors can be categorized into two distinct phases. First, due to the inherent dependency of computational algorithms on numerical representation, raw sequences are transformed into statistical vectors. Second, the statistical vectors are used to train a machine learning or deep learning classifier for the prediction of NT modification sites. Liu et al. [1] developed the very first computational predictor, GPS-YNO2, that discretize protein sequences using precomputed substitution information (BLOSUM62) of the amino acids, and GPS 3.0 (Group-based Prediction System) algorithm that uses vector similarity based scoring strategy for classification. Xu et al. [2] developed another predictor iNitro-Tyr that encodes raw protein sequences into statistical vectors using class aware Position-specific Dipeptide Propensity (PSDP) encoder capable of capturing the occurrence frequencies of Dipeptides in the positive and negative classes at every position in the sequence. Furthermore, they perform classification using a discriminant function approach, that utilizes similarity scores of positive and negative classes. Another predictor, NTyroSite developed by Hasan et al. [3], employs composition of profile-based k-spaced amino acid pairs (pbCKSAAP) as the sequence encoding method. Further, they utilize Wilcoxon’s ranksum feature selection to generate a comprehensive feature space, and Random Forest classifier to discriminate protein sequences between the NT modification and non-modification classes. Nilamyani et al. [4] developed PredNTS predictor, which utilizes four different sequence encoders, namely, K-mer frequency composition, composition of kspaced amino acid pairs (CKSAAP), AAindex and Binary encoding, followed by selection of important features using recursive feature elimination (RFE), four random forest classifiers to generate classification scores from each encoding, and a weighted combination of the four scores to generate the final classification score. More recently, another predictor named PredNitro, developed by Rahman et. al. [7] made use of encoding method similar to class aware Position-specific Dipeptide Propensity (PSDP) encoder, and employs a Support Vector Machine based classifier. The first deep learning based approach, DeepNitro, developed by Xie et al. [5], integrates four encodings of the protein sequences - One-hot encoding (OHE), physiochemical Property Factor Representation (PFR), k-Space spectrum encoding, and Position-specific Scoring Matrix (PSSM) - into a single feature vector, which is then passed to a deep Multi Layer Perceptron (MLP) for classification. iNitroY-Deep predictor developed by Naseer et al. [6], uses integer encoding of amino acids in the protein sequences and Convolutional Neural Network (CNN) based classifier.

A critical analysis reveals that existing predictors have limited performance and are more biased towards detection of false positives, mainly due to the utilization of biased or substandard statistical vectors. Existing predictors make use of diverse types of encoding methods that fall under two main categories, namely, Physiochemical-property and amino acids distribution based. Encoders that belong to the Physiochemical-property class utilize various precomputed values of amino acids for capturing diverse types of sequence information. However, they often fail to extract amino acids sequence order and composition information and show extreme bias towards false positive detection. Different types of encoders that belongs to amino acids distribution, Such as one hot vector encoding based method do not capture correlations of amino acids. Similarly, Position-specific Dipeptide Propensity encoder generates statistical representation by computing class aware occurrence of amino acids and maps train set distribution on test set. These types of encoders rely on distribution of amino acids in train and test sets. They do not perform better when train set has less number of samples and also when distribution of test set slightly differs from train set.

In this study, we propose NTpred, a robust and precise machine learning framework for computational identification of Tyrosine nitration modification sites in protein sequences. we utilized four different composition-based encoding methods for transforming raw protein sequences into statistical feature space. To benefit from diverse types of information captured by different types of encoders, we generate four additional feature spaces by fusing feature spaces of four individual encoders. Furthermore, We eradicate irrelevant and redundant features from each of the eight feature spaces through a recursive feature elimination (RFE) process. At classification stage, we develop a stacked ensemble learning predictor that utilize Gradient Boosted Tree (GBT) classifiers. We generate probabilistic feature space by training eight stacked ensemble learning predictors using eight feature spaces. Generated probabilistic feature space is utilize to train Logistic Regression classifier that makes final predictions of Tyrosine nitration modification and non modification sites.

## Materials and methods

In this section, comprehensive details about the workflow of proposed framework, datasets and evaluation measures are provided. Figure 2 demonstrates 3 core modules of proposed NTpred framework. The *Dataset construction* module illustrates the development process of two benchmark datasets. The *Feature Engineering* module consists of three sub-modules namely Sequence Encoding, Feature Fusion and Feature Selection. Sequence Encoding sub-module employs 4 different encoding methods competent in generating statistical representations of raw protein sequences. With an aim to reap the benefits of different encoding methods, Feature Fusion module produces 4 additional feature spaces by performing early fusion of statistical vectors generated through 4 different encoders. The Feature selection module generates more comprehensive feature spaces by eliminating irrelevant and redundant features from individual encodings and fused statistical vectors. The *Hybrid Ensemble Classifier* consists of a two-stage ensemble classifier, where in first stage, probabilistic feature space is generated by separately passing individual and fused statistical vectors to 8 gradient boosted tree classifiers. In second stage, the probabilistic feature space is used to train a Logistic regression classifier that performs final prediction of modification and non-modification sites. Following sub-sections discusses in-depth details of each of the aforementioned modules.

**Fig. 2.**
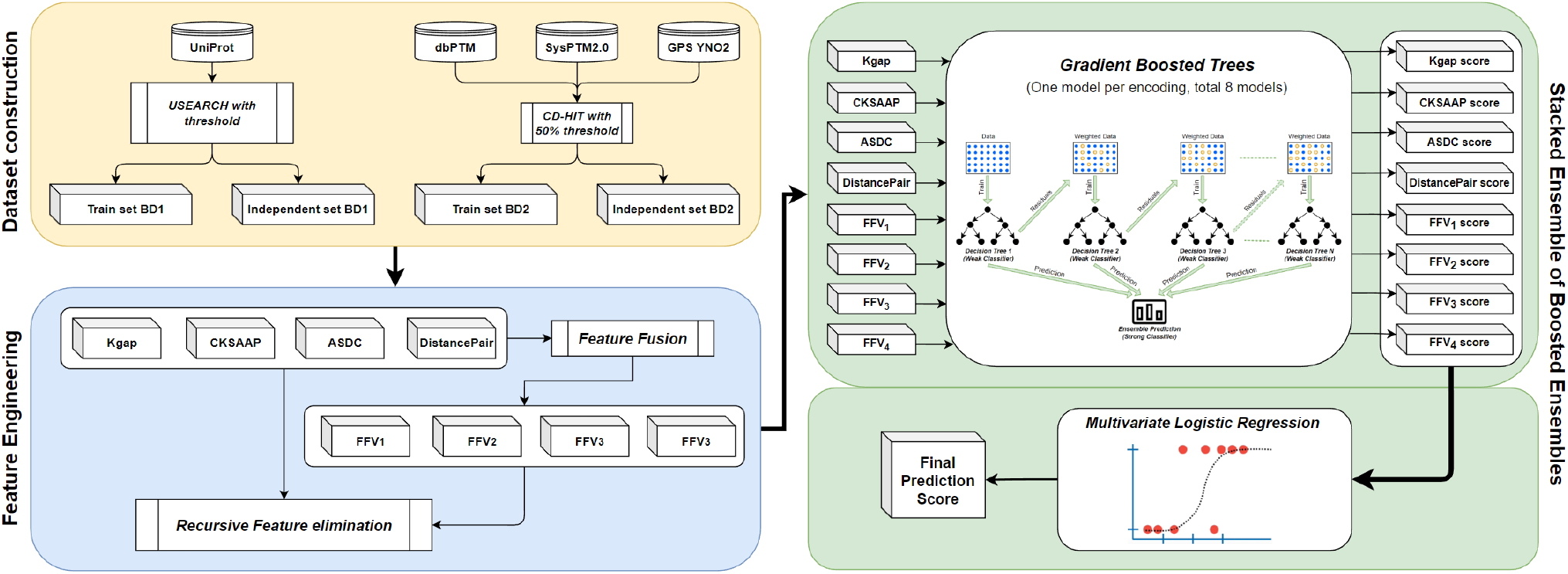
Graphical representation of proposed NTpred framework.

### Numerical representations of protein sequences

To generate numerical representations of protein sequences, the paper in hand makes use of four composition based sequence encoding methods. The following subsections briefly describe the working paradigm of these encoding methods.

#### K-spaced Composition Frequency (Kgap)

Ghandi et al [19] proposed Kgap sequence encoder that captures comprehensive distributional information of amino acids, by considering different gaps among bi-mers in the protein sequences. This encoder has been widely used in diverse types of DNA, RNA and protein sequence analysis tasks such as DNA regulatory sequence identification [20], RNA Pseudouridine sites detection [21], DNA Methylcytosine prediction [22, 24], and small non-coding RNA classification [23].

The encoding process of Kgap encoder can be categorized into two distinct phases. First phase generates 400 bi-mers by utilizing all possible combinations of 20 unique amino acids (20^2^ = 400). Second phase computes distributional information of the generated bi-mers. Suppose a random bi-mer containing X and Y amino acids that appears with different gaps in the sequence as XY, X*Y, X**Y, X***Y, and so on. In the sequence, * denotes a gap containing any random amino acid. The frequency of a bi-mer at various gap levels in a protein sequence is recorded by using gaps of differing lengths.

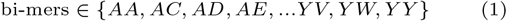

Equation 1 represents the generated set of all 400 possible bi-mers and equation 2 illustrates mathematical expression to compute gap based occurrence frequencies of bi-mers as:

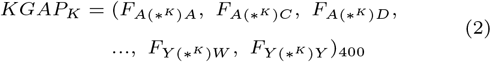

In equation 2, F represents the occurrence frequency of a bi-mer, and (*^*K*^) represents the gap of length K in between the two amino acids in the bi-mer. To clearly demonstrate Kgap sequence encoder process, bi-mers generated at first stage are provided in equation 1, and the rows of Figure 3 describe a hypothetical protein sequence and bi-mers distribution at different gap values. In first row, the parameter K=0 illustrates the occurrence frequencies of bi-mers without any gaps. For example, in the sample sequence, the bi-mer KD occurs once, hence *F_KD_* = 1, PE occurs twice, hence *F_PE_* = 2, bi-mer AA never occurs, hence *F_AA_* = 0, and so on for all the 400 bi-mers. When parameter *K* > 0 is set, gaps are introduced in the bimers and value of K represents length of the gap. In second row, the parameter K=1 illustrates occurrence frequencies of bi-mers with a gap of 1 between the two amino acids in each bi-mer. In the sample sequence, 1-gap-based bi-mer A*P occurs twice as ACP and AYP, hence *F_A*P_* = 2. Similarly, E*T occurs twice as EGT and EPT, hence *F_E*T_* = 2, and so on for all 400 1-gap-based bi-mers. In third row, the parameter K=2 illustrates occurrence frequencies of bi-mers with a gap of 2 in-between the bi-mers. In the sample sequence, 2-gap-based bi-mer A**E occurs twice as ACPE and AYPE, hence *F_A**P_* = 2, T**L occurs once as TYIL, hence *F_T**L_* = 1, and so on for all 400 2-gap-based bi-mers. A similar process is used to compute the occurrence frequencies of bi-mers for higher values of K. Here, we utilize K in range 0 to 4 to generate the complete Kgap feature vector for each sequence. Since each value of K generates a 400 dimensional statistical vector, concatenating vectors from all 5 values of K generates a 2000 dimensional statistical vector.

**Fig. 3.**
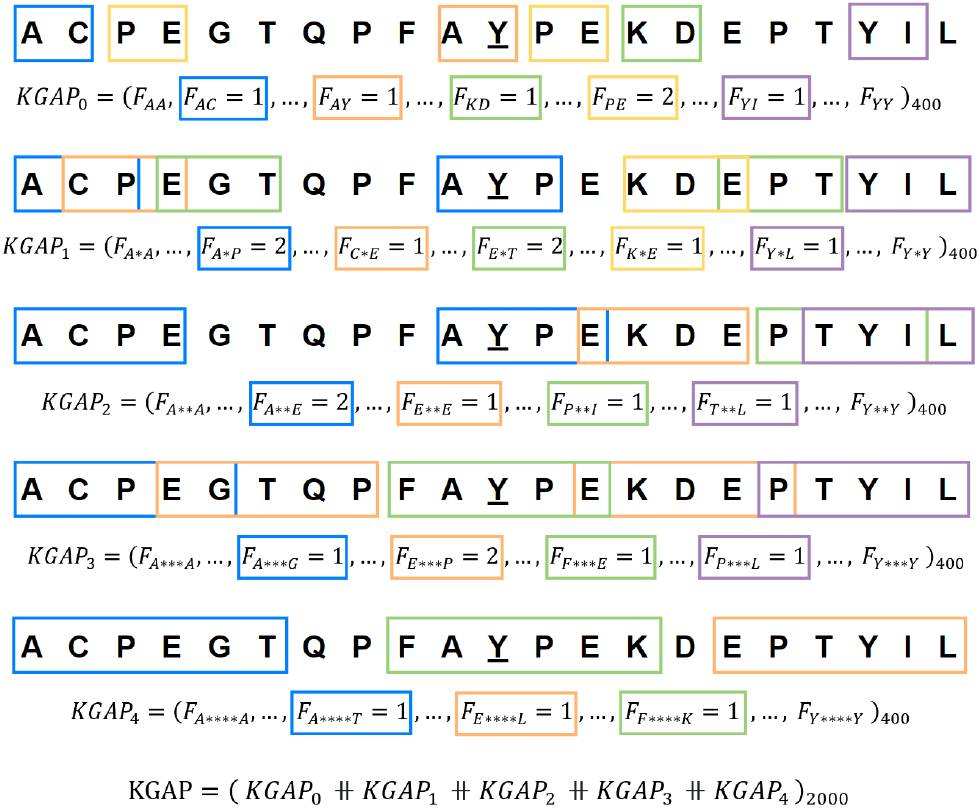
Illustration of the Kgap encoding method.

#### Composition of K-spaced Amino Acid Pairs (CKSAAP)

The CKSAAP sequence encoder, developed by Chen et al. [25], is an extended version of the Kgap encoder discussed in section 2.1.1. The Kgap encoder computes absolute occurrence frequencies of bi-mers. However, the length of protein sequences can vary based on the location of the extracted potential interaction site in the long protein chain, more details in section 2.4. In Kgap encoder, the high variability of sequence length creates bias in the absolute occurrence frequencies of the bimers in the two classes. To mitigate this inherent bias, Chen et al. [25] introduces a sequence length based normalization of the feature vector in the CKSAAP encoder. This encoder has been successfully employed in a variety of protein sequence analysis studies, such as, analysis of protein rigidity and flexibility [25], identification of Protein Phosphorylation and Nitration sites [2, 4, 26], and detection of Protein Pupylation and Succinylation [27, 28].

The working paradigm of this encoder is similar to the two-phase process of Kgap encoder. In addition, it normalises the statistical vectors using sequence length at each gap-length. Adapting the Kgap vector defined in equation 2, the CKSAAP encoded vector can be mathematically formulated as:

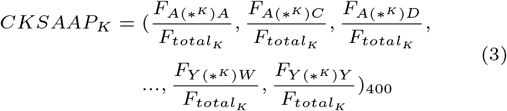

where,

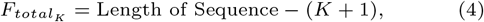

Each row of Figure 4 graphically illustrates the encoding of a hypothetical protein sequence of length 21 into the CKSAAP feature vector. The occurrence frequencies 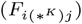 in the numerator are computed exactly like the Kgap encoder. The normalization factor *F_total_K__* varies with K. For K=0, *F*_*total*_0__ = 21 – (0+1) = 20. Similarly, for K=1, *F*_*total*_1__ = 21 – (1 + 1) = 19, and so on for any value of K. In this study, the parameter K in the range 0 to 5 is used to generate the full CKSAAP feature vector. Because each value of K produces a 400-dimensional statistical vector, concatenating vectors from all 6 values of K produces a 2400-dimensional statistical vector.

**Fig. 4.**
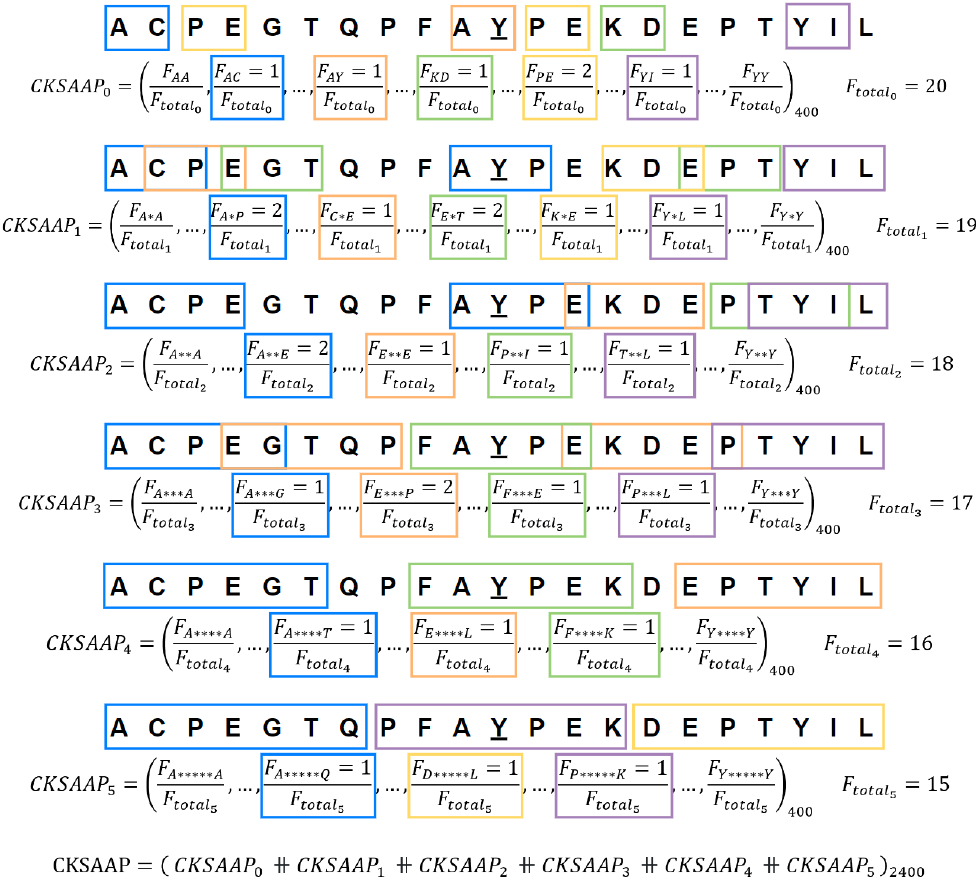
Illustration of the CKSAAP encoding method.

Although normalization potentially removes biases and variations introduced by the genetic sequencing processes and factors such as the length of the extracted sequences, it can adversely affect the biological variability of sequences and s [50, 51]. Furthermore, Lovell et al [52] showed that normalization into the simplex space (where, the components of the vector are only positive and sum to 1) can introduce spurious correlation between the relative compositions even though the absolute compositions are completely uncorrelated. Lovell et al [52] implied that, wherever possible, both absolute abundance of the components and their relative abundance information should be analysed using appropriate compositional features. Hence, in this study, both absolute and normalized composition encoders are utilized.

#### Adaptive Skip Dipeptide Composition (ASDC)

With an aim to control the dimensionality of generated statistical vectors, Wei et. al [29] proposed the ASDC sequence encoder that captures compositional information from all possible adjacent and gap-based bi-mers in a sequence. In both Kgap and CKSAAP encoders, although long range dependencies of amino acids can be captured at large value of K, however it creates sparse high-dimensional feature spaces that adversely affect the performance of supervised learning approaches for classification [49]. To mitigate this issue, Wei et. al [29] proposed to compute the sum of occurrence frequencies of bi-mers at all possible values of K in a sequence. In this way rather than generating 400 * (*K_max_* + 1) dimensional vector, it generates a fixed 400-dimensional statistical vector. Furthermore, to address the bias in absolute occurrence frequencies of the bi-mers based on the length of the sequences, as described in section 2.1.2, normalization is performed. This encoder has been utilized in a variety of protein sequence analysis tasks, such as, to identify cell-penetrating and anticancer peptides in proteins [29, 30, 32], and protein methylation sites [31].

The encoding process of ASDC encoder can be categorized into three distinct phases. In first phase, similar to the previous two encoders, it generates the set of 400 bi-mers. In second phase, it computes the total occurrence frequency of each bimer considering all possible gap-values (K) that lie in range 0 ≤ K ≤ Sequence_Length-2. In the third phase, the total occurrence frequencies of the bi-mers are normalized to generate percentage composition. The ASDC feature vector can be mathematically defined as:

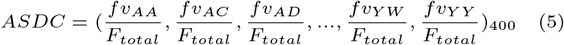

In equation 5, combined occurrence frequency of each bi-mer at all possible K values can be computed as:

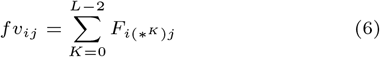

where, 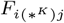 represents the occurrence frequency of bi-mer with length of gap K between *i^th^* and *j^th^* amino acids. The normalization factor in the ASDC feature vector is formulated as:

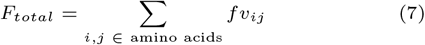

Figure 5 graphically describes the computation of total occurrence frequency of one bi-mer in the ASDC feature vector from a hypothetical sample sequence. The bi-mer AE appears in the sequence twice with gap K=2, hence *F_A**E_* = 2, once with gap K=5, hence *F_A*****E_* = 1, once with gap K=11, hence 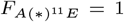, and once with gap K=14, hence 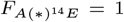. For all other values of gap, it does not occur at all. Hence, in the ASDC encoded vector, the component *fv_AE_* = 2+1 + 1 + 1 = 5. Similarly, the total frequencies are computed for all 400 bi-mers. The 400-dimensional vector is normalized by the sum of all *fv_ij_* components in the ASDC feature vector.

**Fig. 5.**
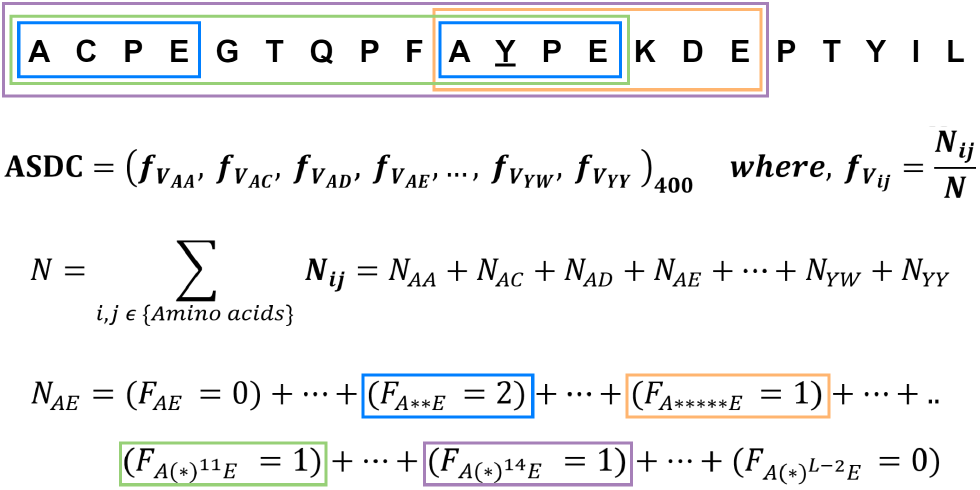
Illustration of the ASDC encoding method.

#### PseAAC of distance-pair (DistancePair)

The Kgap, CKSAAP and ASDC encoders capture distribution information by considering bi-mers with different gap values. However, they do not capture composition of monomers. To address this shortcoming, Liu et al. [33] proposed DistancePair encoder that can capture comprehensive composition information of both monomers and gap-based bimers. This encoder is widely used in DNA and protein sequence analysis studies, such as identification of DNA-binding proteins [33, 34] and anti-cancer peptides [35].

The working paradigm of this encoder can be categorized into three distinct phases. In first phase, the set of all 20 monomers are considered as:

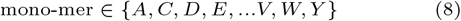

Similar to the previous three encoders, the set of 400 bimers is are generated. In second phase, the occurrence frequencies of monomers and gap-based bi-mers at multiple gap-lengths in the sequence are computed. In third phase, the occurrence frequencies are normalized to encode the percentage composition. For any protein sequence, the DistancePair feature vector can be mathematically formulated as:

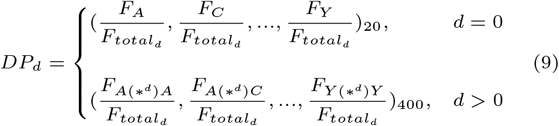

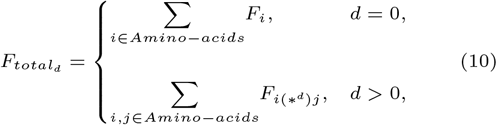

In equations 9 and 10, the parameter *d* controls the level of encoding, *F_i_* represents the occurrence frequencies of monomers, and 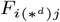 represents the occurrence frequencies of bi-mers with (d-1) random intervening s in between the two amino acids in the bi-mer.

The rows of Figure 6 graphically describes the encoding of a hypothetical protein sequence using DistancePair encoder. In the first row, the parameter d=0 is set that generates the *DP*_0_ vector, which encodes the occurrence frequencies of 20 monomers. For example, amino acid A occurs twice, hence *F_A_* = 2, amino acid P occurs four times, hence *F_P_* = 4, and so on. For any value of *d* > 0, occurrence frequencies of bimers are computed exactly like the Kgap encoder, i.e., d=1 computes occurrence frequencies of bi-mers without any gaps, d=2 computes frequency for bi-mers with a gap of 1 between the two amino acids, and so on. Each *DP_d_* vector is normalized by sum of components of the respective vector, that converts the absolute occurrence frequencies into percentage composition. In this study, the parameter *d* in range 0 to 6 is utilized to generate DistancePair feature vector for each sequence. Since d=0 generates a 20-dimensional statistical vector, and each value of *d* > 0 generates a 400 dimensional statistical vector, concatenating vectors from all 7 values of *d* generates a 20 + (400 * 6) = 2420-dimensional statistical vector.

**Fig. 6.**
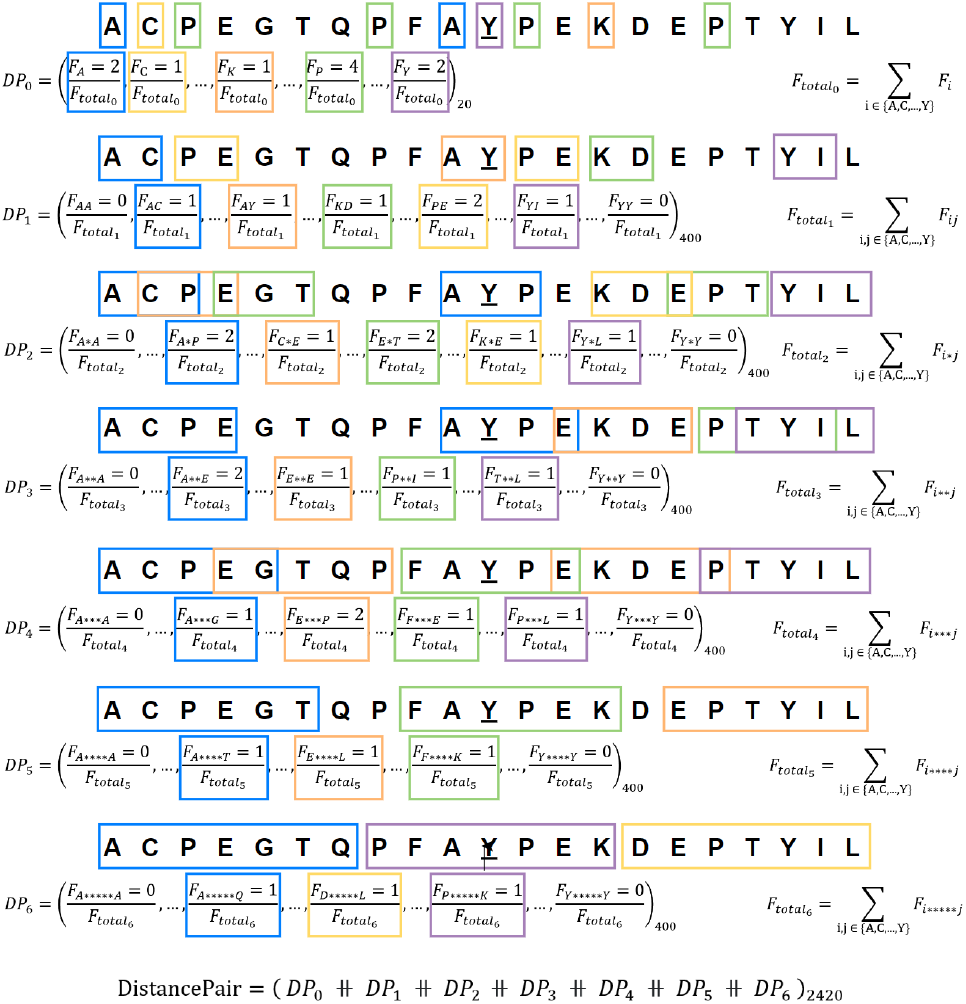
Illustration of the DistancePair encoding method.

### Feature fusion and selection

*Feature fusion* is the process of combining data from several contexts into a single entity that can enhance discriminative information and improves the performance of a computational model. Various feature fusion approaches have been utilized to improve the performance of diverse sequence analysis tasks such as protein sub-cellular localization prediction[36], RNA-protein interaction prediction[37], DNA-binding protein identification [39], and prediction of novel RNA transcripts [38].

To reap the combined benefits of four different encoders discussed in Section 2.1, a vector concatenation based early fusion approach is utilized in our proposed framework. Consider 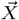 and 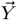 are two different N and M dimensional statistical vectors of a sequence S as:

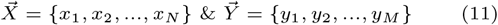

Then, the fused feature vector can be formulated as:

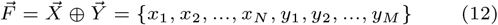

where, 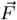 is an (N+M)-dimensional statistical vector of sequence S. Following similar criteria, we create four fused feature vectors as follows:

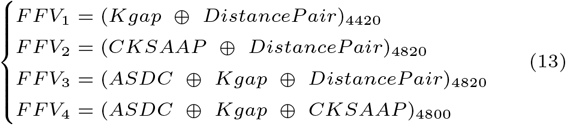

In total, we generate 8 different numerical feature vectors (4 individual sequence encodings and 4 fused feature vectors) for every protein sequence.

*Feature selection* is the process of removing redundant, irrelevant and noisy features that can degrade the performance of predictors. Various feature selection approaches have been utilized in diverse Protein sequence analysis studies, such as, identification of Phosphorylation sites [26], Pupylation sites [27], Succinylation sites [28], cell-penetrating proteins [30], Methylation sites [31], anti-cancer proteins [32] and DNA binding proteins [39]. In the identification of Tyrosine nitration sites also, multiple studies have demonstrated the importance of feature selection methods to build robust computational models. NTyroSite [3] employs a non-parametric feature selection method, Wilcoxon rank-sum (WR) test and PredNTS [4] makes use of the Recursive Feature Elimination process for optimal feature subset selection.

To select important features, we utilize Recursive Feature Elimination (RFE) [40] method with backward step-wise iterative feature selection process. In RFE, a classifier is trained using full feature space, and the features are ordered based on a ranking criterion. In the subsequent iterations, the N lowest ranking features are pruned from the previously ordered features, and again a classifier is trained using reduced feature set. The iteration is terminated when N is greater than or equal to a predefined number of features left. In this study, an Extreme Gradient Boosted Tree classifier along with information gain of the features are utilized as the ranking criterion, with N=50. The RFE process is executed separately for each of the 4 individual sequence encodings and 4 fused feature vectors. The number of features selected for each feature vector, as the stopping criteria for RFE, has been manually optimized by exploring all possible values in the range of 0 to number of dimensions of the feature vector, with step 50, and incorporating them into the proposed framework. After elimination of irrelevant and redundant features, we obtain 950 dimensional feature vector for Kgap, 600 for CKSAAP, 400 for ASDC, 600 for DistancePair, 450 for *FFV*_1_, 600 for *FFV*_2_, 450 for *FFV*_3_ and 700 for *FFV*_4_. For 5-fold evaluation of the model, 80% of the training datasets are used in the RFE process, whereas the complete training datasets are used to perform RFE during the independent testing.

### Hybrid Ensemble Classifier

Ensemble learning is a meta learning process where goal is to improve predictive performance by consolidation of diverse types of models. The working paradigm of ensemble learning approaches can be categorized into two stages. In first stage, utilizing data from a variety of contexts, numerous *weak learners* are trained. In second stage, outputs of the weak learners are intelligently combined to generate the final output. In this way, multiple weak learners together develop a strong learner that demonstrates enhanced generalization capabilities, and reduces model bias and variance [41, 53]. Ensemble learning has been widely utilized in a variety of sequence analysis tasks, such as identification of Nitration and Succinylation in proteins [3, 4, 28], RNA Pseudouridine detection [21], DNA Methylcytosine prediction [22, 24], DNA-binding protein identification [39], prediction of cell-penetrating proteins [29, 30], anti-cancer protein detection [35], and lncRNA-protein interaction prediction [37]. Based on working paradigms, ensemble learning approaches can be broadly categorized into 3 different classes, namely, Bagging, Stacking and Boosting.

*Boosted* ensemble learning [42], is a sequential and iterative process, where multiple weak learners are trained adaptively. An initial weak learner is trained using all observations. Then, the successive weak learners in the sequence are trained iteratively such that the importance of the observations that were misclassified by the previous weak learner are higher than the correctly classified observations. Finally, the predictions from all the weak learners are statistically combined to obtain the final prediction.

One variant of Boosted ensembles is Gradient boosting machines (GBMs) [44–46], that adapts a gradient-descent based formulation of the boosting ensemble method. In GBMs, each new weak learner is iteratively constructed such that they are maximally correlated with the negative gradient of the loss function of the entire ensemble [44], as depicted in Figure 7. With proper choice of the loss function, GBMs are adaptable to both regression and classification tasks. By utilizing Decision Tree classifier with a logistic loss function as the weak learner in a GBM, the *Gradient Boosted Trees* ensemble (GBTs) is obtained [56].

**Fig. 7.**
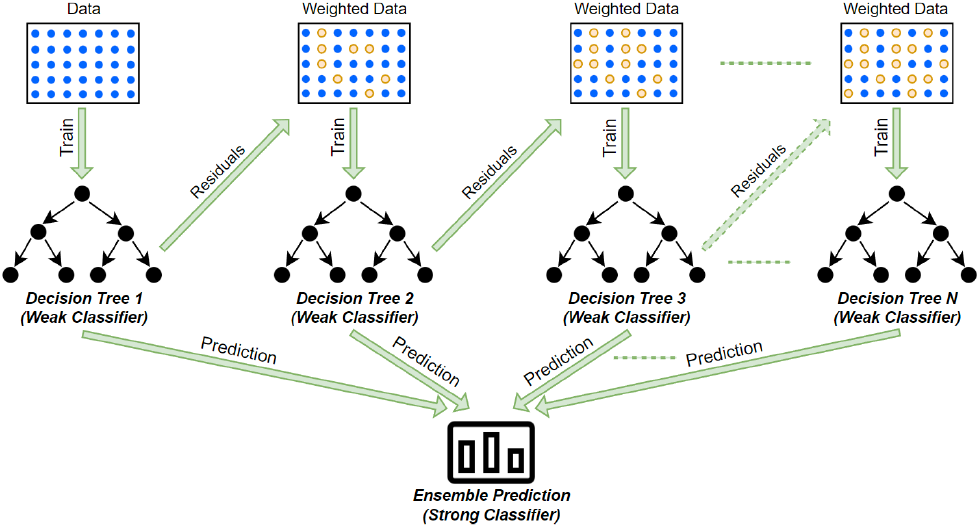
Graphical illustration of Gradient Boosted Tree classifier.

In *Stacked* ensemble learning [43], several weak learners are parallely trained using all observations. Then, a meta learner is trained, using the predictions of the weak learners as the feature space, to generate the final prediction. Here, the motivation behind using a meta-learner is the capability to learn how diverse weak learners perform, identify their benefits, and combine them in the best possible way. An example of a simple meta learner is *Logistic Regression*, which is a multivariate statistical analysis technique, that estimates the probability of occurrence of an event, by fitting a linear combination of the explanatory variables to the logarithm of odds ratio derived from the dichotomous response variable, minimizing the negative log-likelihood (logistic loss) [47].

In this study, a hybrid *Stacked ensemble of Boosted ensembles* architecture has been developed as the classifier model, described graphically in Figure 8. As the weak learners of the stacked ensemble, Gradient Boosted Tree ensembles are trained using each of the eight feature vectors that underwent RFE, as discussed in section 2.2. Then, the predicted probabilities from the eight weak learners are combined to generate a new feature space. Finally, a Logistic Regression model with a linear kernel is trained on this 8-dimensional probabilistic feature space to generate the classification model for final prediction. This final regression step of the hybrid architecture can be viewed as a fusion of information learnt from different compositional context of amino acids in a sequence.

**Fig. 8.**
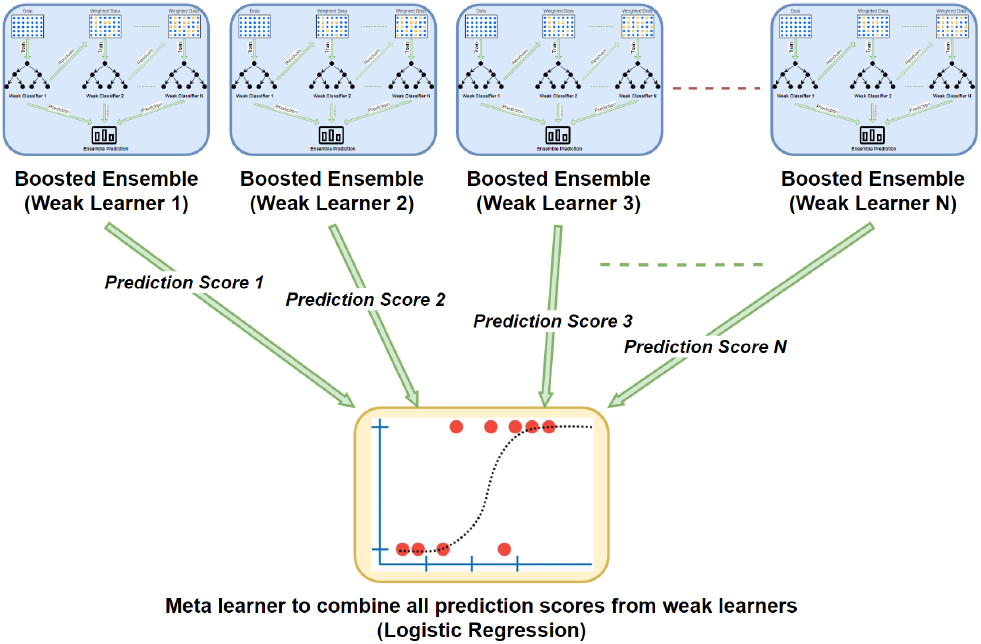
Graphical illustration of Stacked Ensemble.

### Benchmark datasets

To develop a reliable and accurate predictor for Tyrosine nitration modification, stringent selection of suitable datasets to train and test the predictor is a fundamental requirement [2, 3]. In literature, we found two benchmark datasets BD1 and BD2, for Tyrosine nitration site identification. Naseer et al. [6] developed the BD1 dataset by extracting raw protein sequences and their associated NT modification information from the UniProt [69] database. To ensure the quality of data on which classifiers can train in a comprehensive way, by avoiding overfitting and under-fitting, they utilised USEARCH algorithm [54] with a 70% similarity threshold to remove homologous sequences. Nilamyani et al. [4] developed the BD2 benchmark dataset by utilizing annotated data from the dbPTM [67], SysPTM2.0 [68] and GPS-YNO2 [1] databases. They discarded redundant sequences by utilizing CD-HIT [55] with a similarity threshold of 40%.

Both authors have provided the benchmark training datasets, along with the independent test sets. Figure 9 illustrates the distribution of samples in the training and test sets for both datasets. BD1 dataset contains 1191 samples of modification class (positives) and 1191 samples of non-modification class (negatives) in the train set. In the independent test set, it contains 203 positive and 1022 negative samples. Similarly, the BD2 dataset contains 229 positive and 354 negative samples in the train set, and 98 positive and 153 negative samples in the independent test set.

**Fig. 9.**
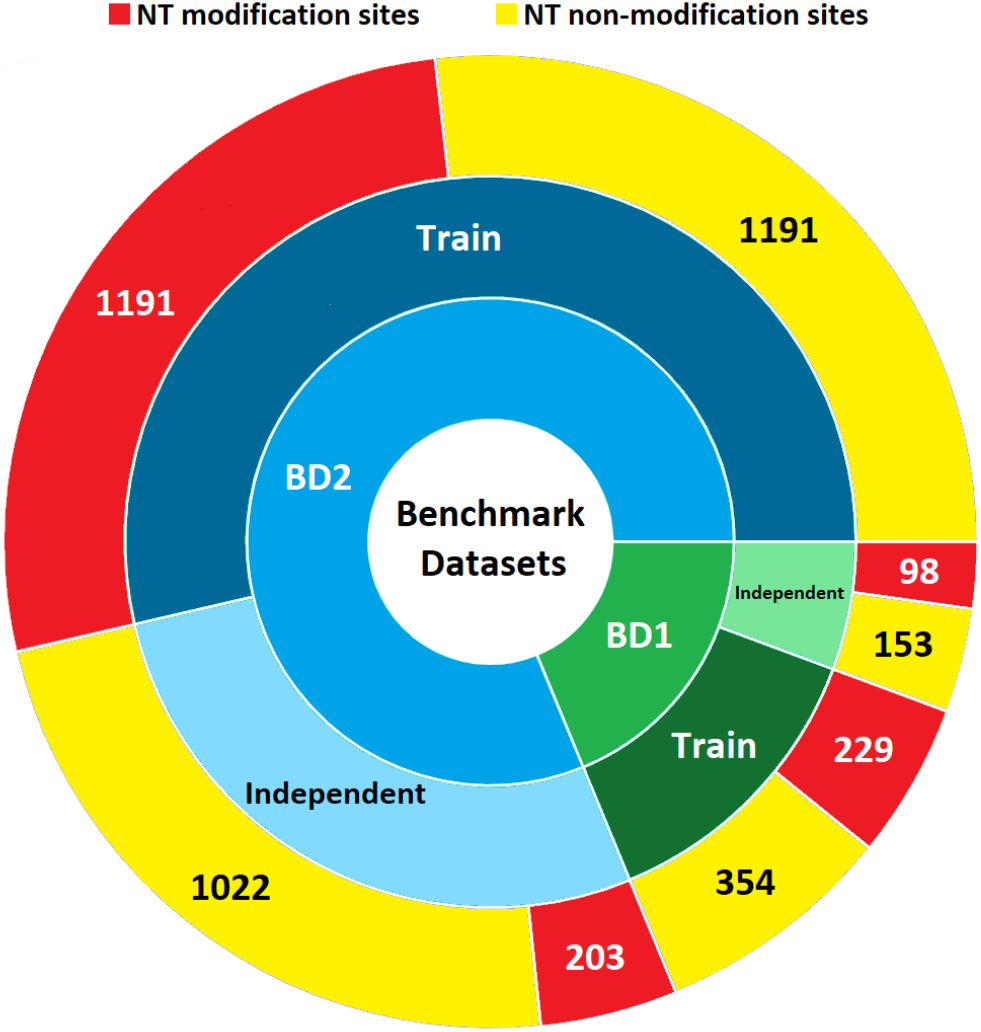
Statistics of benchmark Datasets.

In the construction of both BD1 and BD2 benchmark datasets, positive and negative sequences are extracted from long protein chains using a window represented in Chou’s scheme [15] that contains the potential Tyrosine modification site at center of the sequence as:

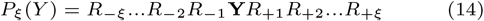

In equation 14, *R_i_* represents the flanking region around the potential modification amino acid residue in the sequence, with a window size of *ξ*, as illustrated in Figure 10. By sliding this window along every long protein chain, sequences containing Y (tyrosine) at the center are extracted. Sequences extracted from experimentally verified modification locations are labelled as positive, and all others are labelled as negative. Both datasets utilize a window size *ξ* = 20. However, since Tyrosine can also occur at either ends of a long protein chain (for example, in Figure 10, locations closer to N and C), these segments extracted using the sliding window can contain missing sections of the sequence at either end. Due to this, the extracted sequences vary in length from *ξ* + 1=21 to 2*ξ* + 1=41. Furthermore, the relative position of each amino acid residue in the segment, according to the window, are maintained by representing missing residues with dummy residue X. But these dummy residues are discarded during the numerical encoding of sequences in our proposed framework.

**Fig. 10.**
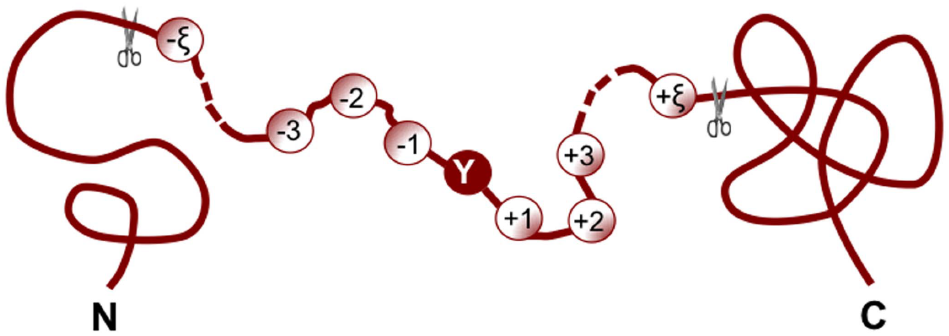
An illustration of Chou’s scheme for a protein sequence containing 2ξ + 1 s with tyrosine (Y) at the center. Adapted from Xu et. al [2] under CC BY 4.0 license.

### Evaluation criteria and performance measures

Following the evaluation criteria of existing predictors [1–7], the proposed NTpred framework is evaluated in two distinct settings. In the first setting, the framework is evaluated on two benchmark training datasets through 5-fold cross validation, which provides an estimate of the predictor’s performance, generalizability and stability with differing training sample sets. In this setting, the training datasets are randomly partitioned into 5 stratified subsets, which maintain the original class ratio. Then, in each iteration, a discreet subset is selected as the evaluation set, whereas the remaining four subsets are combined to train the model. In the second setting, the framework is trained using the entire benchmark training datasets. Then, the respective independent test sets are used to generate performance metrics for the framework. The independent test sets are also employed, either to generate performance metrics for existing predictors of Tyrosine nitration, or utilize the performance information already available in literature. This enables appropriate comparison of proposed framework’s performance with existing predictors.

Furthermore, performance evaluation of the proposed NTpred framework is performed using 4 different measures, namely, Sensitivity (Sn), Specificity (Sp), Accuracy (Acc) and Matthews Correlation Coefficient (MCC). These measures makes use of different factors to compute the performance score. The factors are illustrated in Table 1, where *T_p_, T_n_, F_p_* and *F_n_* are the counts of True Positives, True Negatives, False Positives and False Negatives respectively. The statistical measures can be formulated as:

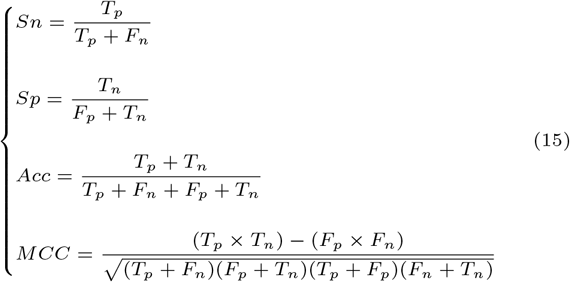

**Table 1.**
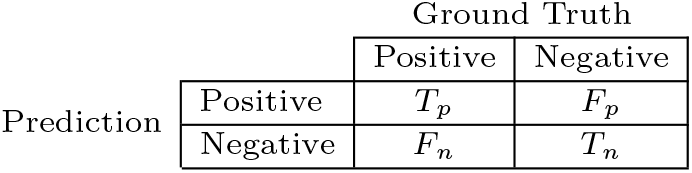
Confusion Matrix

Additionally, the Receiver Operating Characteristic curve (ROC), which plots the *Sn* vs. 1 – *Sp* in the entire range of possible classifier scores, is also analyzed. The Area under the ROC (AUC) is equal to the probability that the classifier will score a randomly chosen positive sample higher than a randomly chosen negative sample irrespective of the class distribution [48]. The AUC can be used to determine optimal classification score thresholds, and measure the overall robustness of the classifier.

## Experimental setup

The proposed NTpred approach is developed in Python, where, the numerical representations are generated by utilizing 2 APIs: *MathFeature* [72] and *iLearnPlus* [73], and the ensemble classifier is developed using 2 APIs: *scikit-learn* [75] and *XGBoost* [74]. Intrinsic analysis of amino acid’s distribution in protein sequences is performed using *Logomaker* [76]. Additionally, by optimizing the hyper-parameters of Machine Learning algorithms, their performance can be improved and regularized. For example, for Gradient Boosted Trees, utilizing 3 optimal hyper-parameters namely, *Number of Estimators* which controls the number of trees in the ensemble, *Maximum Depth* which controls growth of each tree by limiting maximum decision nodes before a final decision is arrived, and *Learning Rate* controls the proportion of residuals (loss) which is used to update weights of samples utilized to train consecutive trees in the ensemble. Furthermore, both Gradient Boosted Trees and Logistic Regression can be further optimized using Lasso (L1) [60] and Ridge (L2) [61] regularization, that penalizes unnecessary complexity of models, promotes relevant features and avoids overfitting [59]. L1 regularization controls absolute weights of model parameters, whereas L2 regularization controls the squared weights of model parameters [59].

Hence, a grid search process [71] is employed to optimize hyper-parameters of Gradient Boosted Trees and classifiers. Table 2 describes values utilized for each hyper-parameter in the grid, and their optimal value identified through the search process. Additionally, all possible combinations of the 4 numerical representations for feature fusion are also explored. Furthermore, to train all 8 Gradient Boosted Tree based classifiers, an early stopping criteria is utilized by monitoring the logistic loss of prediction. Both classifiers, GBT and Logistic regression, are trained with class weights to control class bias of the models.

**Table 2.** Hyper-parameter optimization using grid search.

| Hyperparameters | Grid Search<br>Hyperparameter space | Optimal Hyperparameters |  |
| --- | --- | --- | --- |
|  |  | Gradient Boosted Trees | Logistic Regression |
| *No. of Estimators* | 10, 50, 100, 500, 1000 | 100 | - |
| *Max Depth* | 1, 3, 5, 7, 10 | 3 | - |
| *Learning Rate* | 0.0001, 0.001, 0.01, 0.1 | 0.1 | - |
| *reg_alpha* | 0, 0.00001, 0.001, 0.01, 0.1, 1 | 0 | 0 |
| *reg_lambda* | 0, 0.00001, 0.001, 0.01, 0.1, 1 | 1 | 1 |

## Results and Discussions

This section illustrates distribution of amino acids in protein sequences. Over two benchmark datasets, it quantifies predictive performance and generalizability achieved by proposed NTpred framework at various stages. Furthermore, it summarizes intrinsic visual analysis of 8 numerical feature spaces, and performs their comparison with probabilistic feature space generated through predictions of first stage classifiers. Finally, it describes performance comparison of proposed framework with existing Tyrosine nitration predictors.

### Amino-acids distribution analysis

Figure 11 describes discriminative distribution potential of amino-acids in proteins of two different classes for two benchmark datasets. In protein sequences, at each position prevalence of amino acids is depicted through the height of characters that represents amino acids. To reduce clutter that occurs when all amino acid occurrence frequencies at every position is visualized, only those amino acids whose relative abundance is at least 7.5% are plotted. In Figure 11, different colors of characters represent five dominant chemical properties of the amino acids, namely Polar, Neutral, Basic, Acidic and Hydrophobic.

**Fig. 11.**
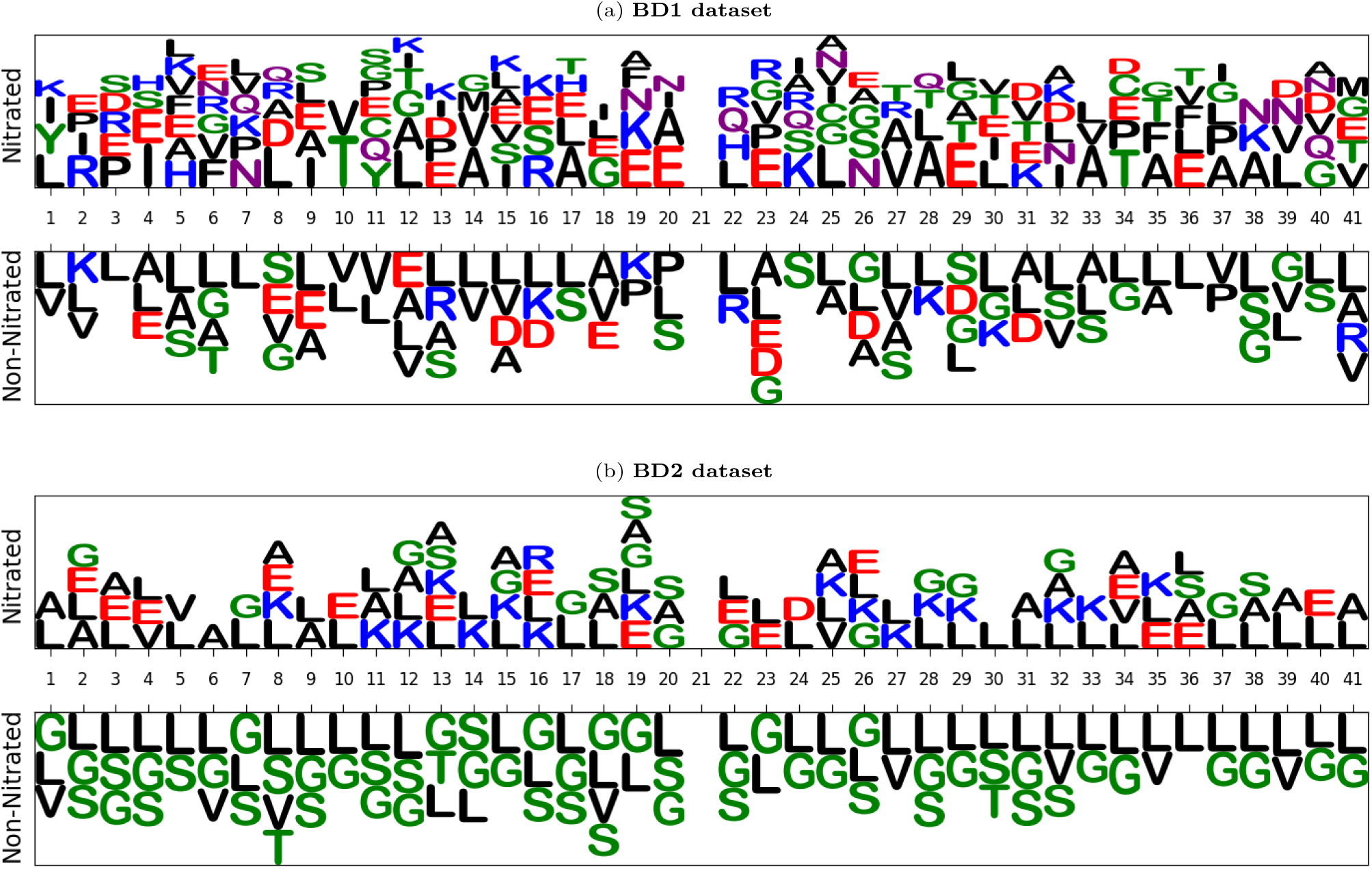
Position-specific distribution analysis of amino acids. Color profile represents the dominant chemical property of the amino acids: *Green-Polar, Purple-Neutral, Blue-Basic, Red-Acidic, Black-Hydrophobic*.

According to intrinsic visual analysis, negative class sequences have a significantly higher prevalence of Hydrophobic and Polar amino acids, whereas positive class sequences have a high prevalence of Neutral, Acidic, and Basic amino acids. Thus, a difference in the amino acids composition between the sequences of positive and negative classes is observable. This discriminative distribution of amino acids suggests that compositional feature encodings of the protein sequences could prove effective to build robust and precise framework for the identification of Tyrosine nitration sites. Furthermore, composition of amino acids in the sequences of two benchmark datasets is noticeably different, specifically in both classes, sequences of BD1 dataset are significantly more diverse than sequences of the BD2 dataset. For example, among positive and negative class sequences, difference in abundance of the most common amino acid Leucine (L) is significantly greater in BD1 dataset than in BD2 dataset.

Additionally, from Figure 11 it can also be concluded that discriminative distribution of amino acids becomes more prominent when we consider occurrence of two consecutive amino acids pairs, as well as amino acids pairs occurrence with particular gap between them. For example, in both datasets amino acid pair LL occurs with considerably higher frequency in negative class than positive class sequences. It is also evident that Hydrophobic and Polar amino acid residues occur more frequently in each other’s neighborhood in positive class sequences than in negative class sequences. The occurrence frequency of the pair LL with small gap of varying lengths in between them is also higher in negative class sequences than in the positive class sequences. Similarly, other amino acids pairs, such as LS and LG, also demonstrate a similar distribution patterns in BD1 and BD2 datasets respectively. The characteristics of amino acids distribution in the sequences reveal that composition based sequence encoding methods are more suitable to transform protein sequences into statistical vectors, as these methods capture diverse types of amino acids distributional information such as neighboring as well as gap-based amino acid pairs.

### In-depth Performance analysis of proposed NTpred framework

Tables 3 and 4 illustrate predictive performance of GBT classifier in terms of 5-fold cross validation and independent test based settings using 4 numerical encodings (Kgap, CKSAAP, ASDC and DistancePair) and 4 fusion based numerical encodings (*FFV*_1_, *FFV*_2_, *FFV*_3_ and *FFV*_4_). Both Tables also show performance of GBT classifier when fed with the optimized subset of features selected through Recursive Feature Elimination (RFE) process. Furthermore, it also presents the predictive performance of our proposed NTpred framework.

**Table 3.**
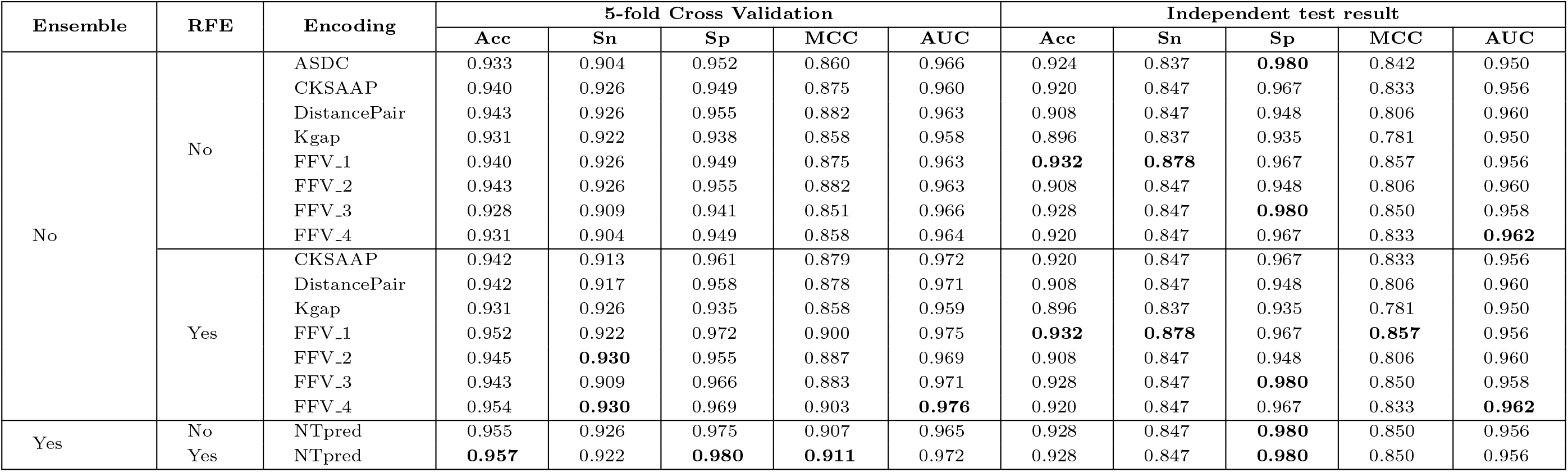
Performance of individual encodings, fusion vectors, and the NTpred framework on the BD1 dataset.

**Table 4.**
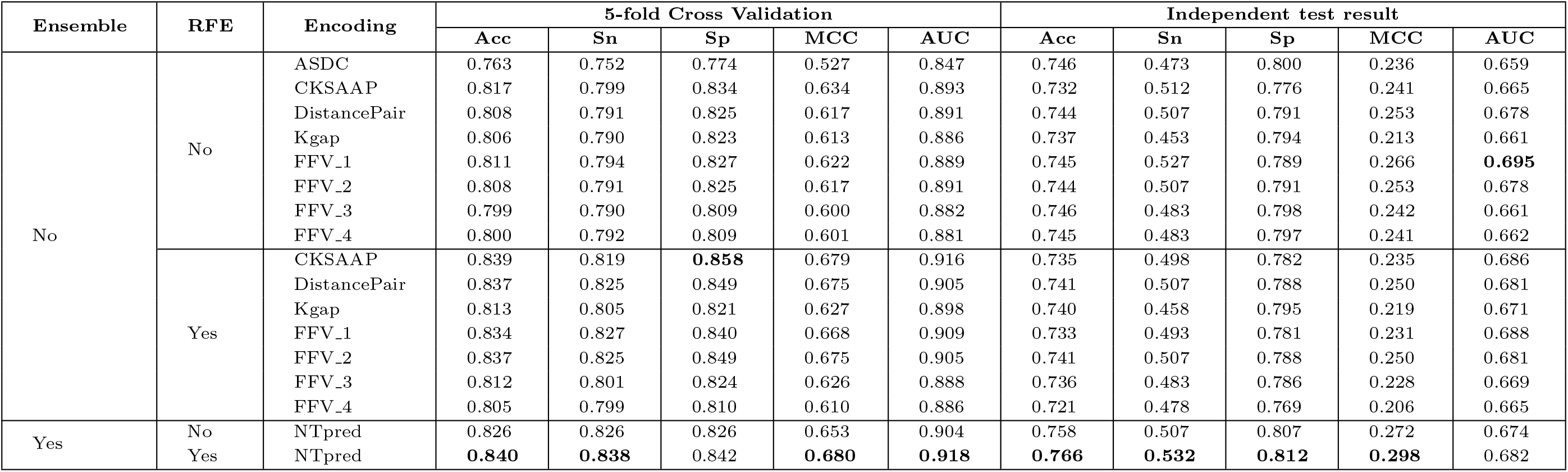
Performance of individual encodings, fusion vectors, and the NTpred framework on the BD2 dataset.

As evident from Table 3 on BD1 dataset in both 5-fold cross validation and independent test based settings, among 4 individual encodings using the full feature space, DistancePair encoder facilitates GBT classifier to produce better performance compared to other encoders. Furthermore, although fused numerical encodings have similar performance compared to individual encodings in 5-fold cross validation setting, however in the independent test setting they outperform the individual encodings with 1% to 2% higher average values across all 5 measures. Among 4 fused numerical encodings, *FFV*_1_ outperforms others in both 5-fold cross validation and independent test settings. Feature selection over the 4 individual encodings have very minute improvement in performance in both settings. However, the 4 fused numerical encodings achieve significant performance boost in 5-fold cross validation, with 1% to 5% higher values across all 5 measures.

Similarly, from Table 4 on BD2 dataset, among the 4 individual encodings using the full feature space, the CKSAAP performs best in 5-fold cross validation, whereas DistancePair performs best in independent test setting. Among the fused numerical encodings, *FFV*_1_ performs best in both settings. Analogous to performance on BD1 dataset, the fused numerical encodings in BD2 dataset have similar performance compared to individual encodings in 5-fold cross validation, but outperform the individual encodings with 1% to 2% higher average values in all 5 measures for the independent test setting. Unlike the BD1 results, after RFE process optimizes the feature space in BD2 dataset, the individual encodings as well as fused numerical encodings show significant improvement in predictive performance with 1% to 6% higher values across all 5 measures in 5-fold cross validation.

Upon exploiting the benefits of all 4 individual encodings and 4 fused numerical encodings in the proposed hybrid ensemble framework of NTpred, significant performance improvements are observed. From Table 3 on BD1 dataset, the NTpred framework acheives 1% to 7% higher values across all 5 measures in comparison with the 8 individual features spaces, in both 5-fold cross validation and independent test settings. This improvement is observable using full feature spaces as well as RFE based optimized feature spaces. Similarly, from Table 4 on BD2 dataset, the proposed framework acheives 1% to 13% higher performance values across all 5 measures in comparison with the 8 individual features spaces, in both 5-fold cross validation and independent test settings. Using RFE optimized feature space, the NTpred framework acheives 1% to 9% higher performance measures in comparison with the 8 individual features spaces in both settings. Additionally, the framework performs better when RFE based feature selection is added to its pipeline. However, this improvement is more evident on BD2 dataset with 1% to 3% improvement in performance in both settings, whereas the improvements are approximately 0.5% on BD1 dataset. One additional observation is that the discrepancy in performance of GBT classifier using individual encodings and the NTpred framework between the BD1 and BD2 datasets can be attributed to the differences in the amino acids distribution between the datasets, as discussed in Section 4.1.

### Intrinsic analysis of feature spaces

To investigate the discriminative potential of 4 individual encoders, 4 fused numerical encodings, and generated probabilistic feature space, we perform an intrinsic visual analysis. Because the feature spaces are significantly highdimensional and sparse, effective visual analysis requires dimensionality reduction in order to construct appropriate human-readable plots. In this analysis, dimensionality reduction is carried out in two stages. In first stage, highdimensional numerical representations are passed to a linear Principal Component Analysis (PCA) [57] and the top 5 components are extracted. In second stage, the reduced 5-dimensional feature spaces are passed to T-distributed Stochastic Neighbor Embedding (t-SNE) [58] that generates a 2-dimensional feature spaces and make visualizations. However, we must keep in mind that this drastic reduction in dimensionality may result in significant loss of information and variance in the data.

Figure 12 graphically visualizes nine different statistical representations of modification (positive) and non-modification (negative) sites for both BD1 and BD2 benchmark training datasets. Each of the 4 statistical vectors generated through compositional encoding methods and 4 fused feature vectors are individually visualized. Similarly, the Recursive Feature Elimination (RFE) based reduced statistical vectors are also individually analyzed. Additionally, the probabilistic feature space generated from the prediction of the 8 Gradient Boosted Trees (GBTs) using the 4 compositional encodings and 4 fused numerical vectors in the first stage of our proposed framework is examined. The probabilistic feature space is generated in a 5-fold approach for both benchmark training datasets, where 4 folds are used to train the GBTs and the remaining fold is used to generate the probabilities.

**Fig. 12.**
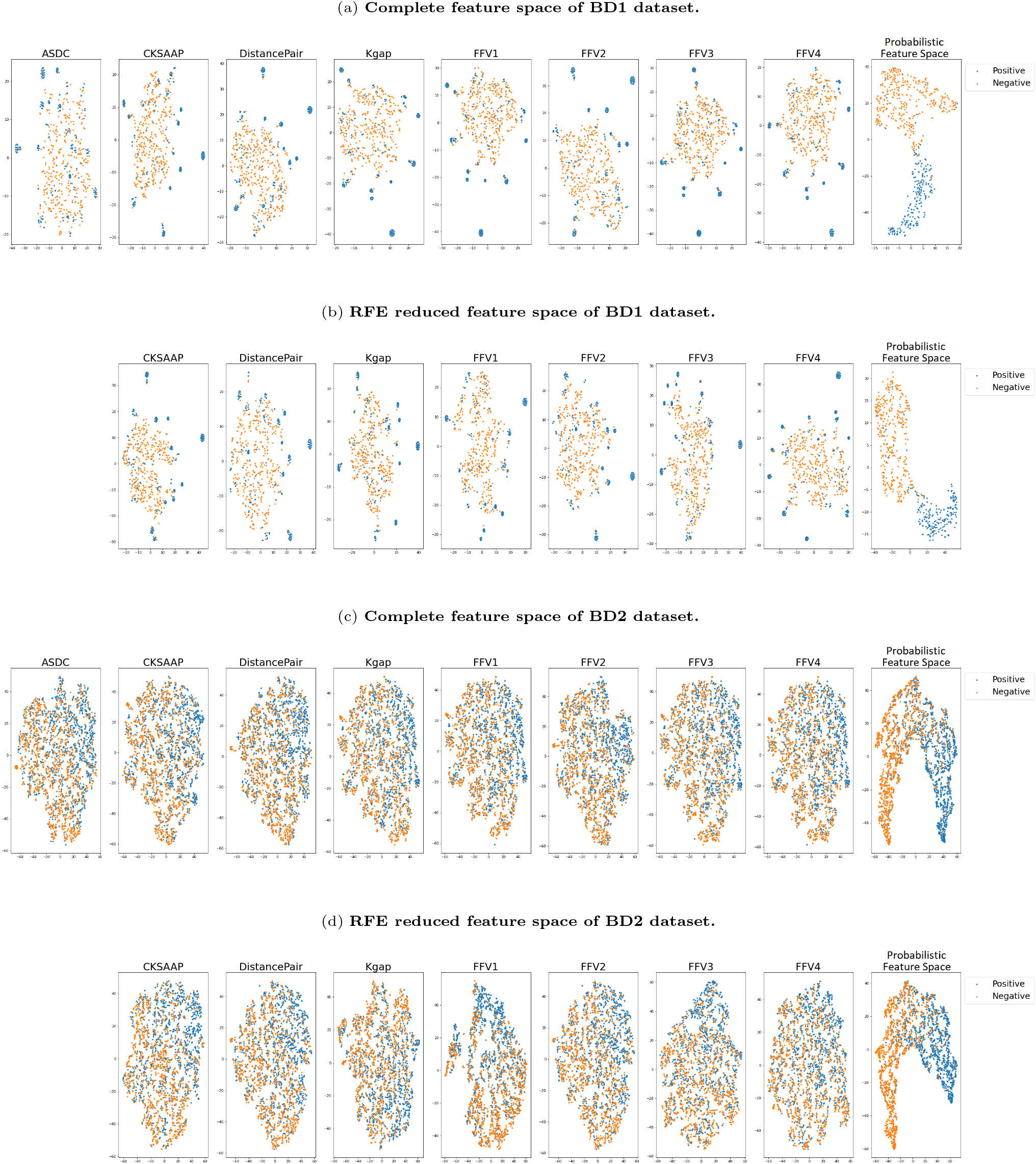
Visual analysis of Numerical encodings, Fusion vectors and generated Probability space. Each high dimensional feature space is separately treated with a combination of PCA and tSNE to map to a 2-dimensional space that can be visualized.

Figure 12a shows 4 individual vectors, 4 fused vectors and generated probabilistic feature space for the complete feature spaces of BD1 dataset. Among individual encoder vectors, ASDC encoder vectors forms more overlapped clusters compared to CKSAAP, DistancePair and KGAP encoder vectors which show slightly better separation between the clusters of the two classes. Furthermore, it is evident that by exploiting combinations of compositional encodings in the four fused feature vectors, all of them show better separation between clusters of the two classes. Similarly, analyzing the Figure 12b on the RFE reduced feature spaces of BD1 dataset reveals almost identical patterns.

In contrast over BD2 dataset, Figure 12b reveals that for all 8 encoded vectors, unlike BD1 dataset, there is almost negligible separation between clusters of positive and negative classes. Almost all encoded vectors form a single large cluster with significantly higher overlap between samples of both classes. However, it is also evident that within the large cluster of all 8 feature vectors, the positive vectors are more densely packed towards one side of the cluster. From Figure 12d that visualizes RFE reduced feature spaces of BD2 dataset, although it demonstrates similar patterns as the complete feature spaces in Figure 12c, the positive vectors are slightly more densely packed in RFE reduced feature spaces than complete feature spaces.

Furthermore, the analysis of all 4 plots in Figure 12 also reveal that the probabilistic feature space generated through GBTs is significantly more discriminative than the 8 numerical encodings. From Figures 12a and 12b on the BD1 dataset, it is evident that two major clusters form, one for each class. The clusters of the complete feature space have good separation, but the clusters themselves are less cohesive. However, the clusters of the RFE reduced feature space show better cohesion also. Similarly, in Figures 12c and 12d for the BD2 dataset, two major clusters are generated. Although these clusters for the BD2 dataset have slight overlap, they are significantly better separated than the 8 numerical encodings where the separation is less prominent.

The analysis reveals that utilizing a two-stage stacked ensemble classifier increases the discriminative potential of our NTpred framework. Additionally, the performance difference of our proposed predictor between the BD1 and BD2 benchmark datasets is also described by the plots, where two classes are significantly more separable in BD1 dataset in comparison to BD2 dataset.

### Comparative analysis of proposed NTpred framework with existing predictors

According to best of our knowledge, to date 7 different computational predictors [1–7] have been developed for the identification of Tyrosine nitration sites in protein sequences. Among them, only iNitroY-Deep [6] predictor is evaluated on BD1 dataset. Five predictors namely GPS-YNO2 [1], DeepNitro [5], NTyroSite [3], PredNTS [4] and PredNitro [7] are evaluated on BD2 dataset. Predictors performance on BD2 dataset is gathered from the published results of PredNTS [4]. The source code of PredNitro [7] predictor is available, so we computed its performance on both benchmark datasets. However, the iNitro-Tyr predictor [2] is not evaluated on BD1 or BD2 datasets. Furthermore, source-code and dataset used in this predictor are not available, hence we exclude it from our comparative analysis.

Figure 13 illustrates performance comparison of proposed framework with two existing predictors on BD1 dataset. As shown in Figure 13a and 13b, among two existing predictors, in terms of both 5-fold cross validation and independent test set based evaluations, PredNitro [7] predictor that made use of Position-specific conditional probability based feature encoding method and Support Vector Machine classifier performs best by producing 93.2% accuracy, 81.2% MCC and 86.7% AUC 5-fold Cross Validation, and 93.1% accuracy, 81.2% MCC and 86.0% AUC in Independent test evaluation. The iNitroY-Deep [6] predictor, that utilized integer encoding of amino acids along with a Convolutional Neural Network based classifier is evaluated only on the independent test set of BD1 dataset, performs worst by achieving 87.2% accuracy, 74% MCC and 91% AUC. Basically, replacing amino acids with random integers generates poor representation of protein sequences because this encoding strategy does not capture any dependencies between different amino acids in the sequences which degrade the performance of iNitroY-Deep predictor.

**Fig. 13.**
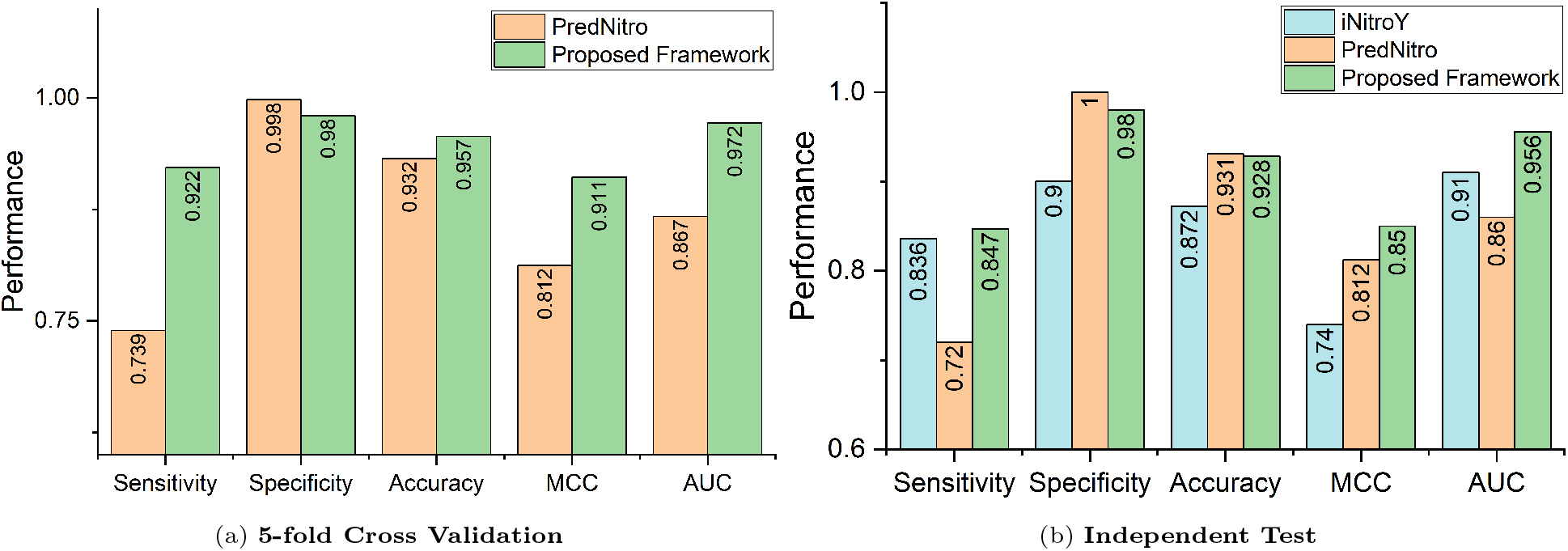
Performance comparison of proposed and existing predictors on BD1 dataset.

In contrast, although proposed NTpred framework performs similar to PredNitro in terms of accuracy in independent test evaluation on BD1 dataset, however, across other two evaluation measures namely MCC and AUC, it outperforms PredNitro by producing 3.8% and 9.4% higher performance respectively. Furthermore, in 5-fold Cross Validation on BD1 dataset, our proposed framework outperforms iNitroY-Deep with 2.5%, 9.9% and 10.5% higher performance in terms of Accuracy, MCC and AUC.

Figure 14 illustrates performance comparison of proposed framework with five existing predictors on BD2 dataset. As shown in Figures 14a and 14b, among existing predictors, in terms of both 5-fold cross validation and independent test set based evaluations, PredNTS predictor [4] that made use of four different encoding methods and Ensemble classifier, produce highest performance by achieving 81.9% accuracy, 63.9% MCC and 91% AUC over 5-fold cross validation, and 76.1% accuracy, 28.6% MCC and 68.0% AUC over independent test set. The NTyroSite predictor [3] that is based on Sequence Evolutionary encoding method and Random Forest classifier, achieved second highest performance in terms of accuracy and MCC. NTyroSite predictor generates statistical representation by utilizing Physiochemical-property based encoding methods that do not capture amino acids correlation information, which eventually degrade the performance of predictor.

**Fig. 14.**
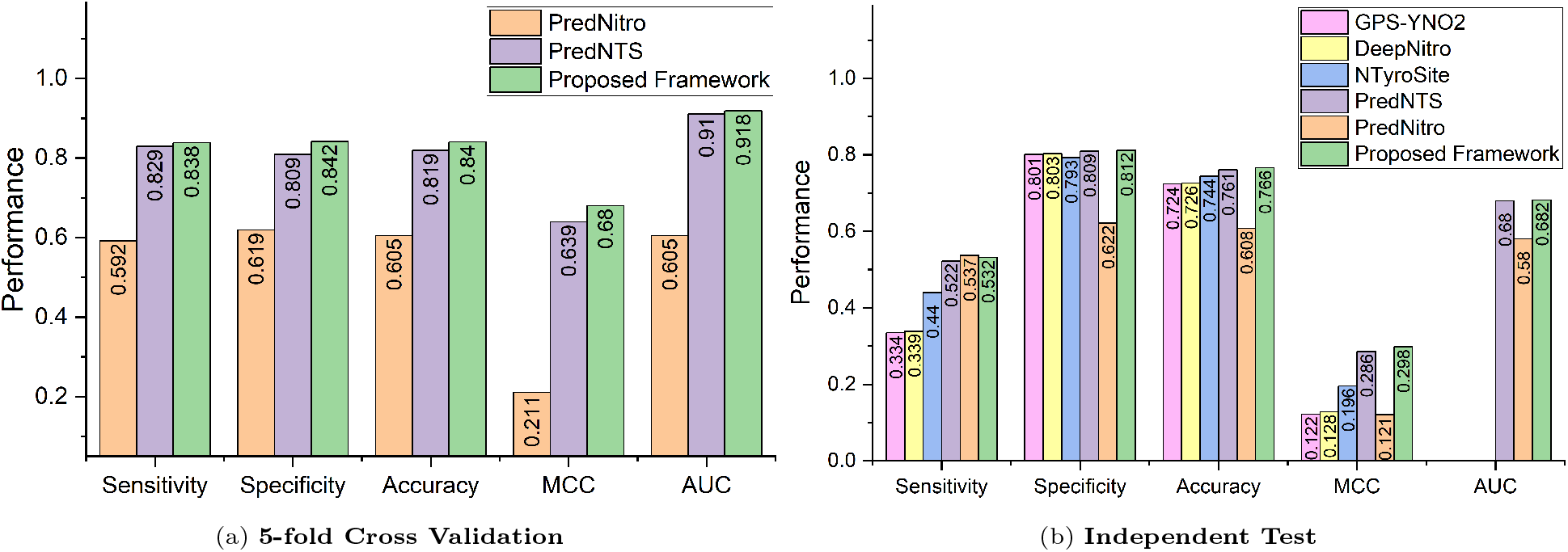
Performance comparison of proposed and existing predictors on BD2 dataset.

The GPS-YNO2 [1] and DeepNitro [5] predictors produce almost similar performance and manage to achieve 3rd rank on the BD2 dataset. This shows that the BLOSUM62 and Position Specific Scoring Matrix (PSSM) based encodings, utilized by GPS-YNO2 and DeepNitro respectively, only compute position specific physiochemical features, and fail to capture sequence level discriminative information. Although PredNitro [7] predictor produced highest performance on BD1 dataset, however it produce lowest performance on BD2 dataset. This performance gap reveals that Position-specific Conditional Probability based encoding method is distribution dependent and only generates better representation if the train and test sets have very similar distributions. It will not produce a better representation if there is a slight difference between the distribution of train and test sets, because it computes distribution of amino acids from the train set and maps them on the test set.

On the other hand, proposed NTpred framework once again produce highest performance on BD2 dataset. Although, proposed and PredNTS predictors produce almost similar performance over Independent test evaluation on BD2 dataset. However, in 5-fold Cross Validation on BD2 dataset, in comparison to PredNTS predictor, the proposed NTpred predictor produces 2.1%, 0.9%, 3.1%, 4.1% and 0.8% higher performance in terms of accuracy, sensitivity, specificity, MCC and AUC. Hence, overall it can be concluded that existing predictors are not generalized, as they produce better performance only on partial datasets. Contrarily, the proposed predictor is more generalized as it produce almost similar performance on both datasets.

## Conclusion

Tyrosine nitration site prediction is important to understand diverse biological processes in various Neurodegenerative, Cardiovascular and Autoimmune diseases, and Carcinogenesis in humans, and support the discovery of novel therapies and drugs. Several computational predictors have been developed for identification of Tyrosine nitration sites, however they lack in performance mainly due to utilization of ineffective sequence encoding methods which do not capture discriminative potential of the amino acids, and introduces bias with disproportionately higher Specificity than Sensitivity. The paper in hand presents a robust computational framework that is competent in precisely identifying Tyrosine nitration sites in proteins. The proposed NTpred framework employs a variety of compositional encoders capable of capturing discriminative information of amino acids in protein sequences. Furthermore, the fusion of features from different encoders and selection of informative features enhances the predictive capability of NTpred framework. A thorough performance analysis of proposed framework reveals that incorporating information from different contexts into the hybrid ensemble architecture significantly improves the predictor’s generalizability, robustness and predictive power. An in-depth comparison of our proposed NTpred framework with six existing predictors using two benchmark datasets demonstrates higher performance potential of proposed framework compared to state-of-the-art. A compelling direction for future studies would be to investigate the effectiveness of various Natural Language Processing techniques, including traditional statistical analysis, language models, and cutting-edge neural network architectures.

## References

1. Liu, Zexian & Cao, Jun & Ma, Qian & Gao, Xinjiao & Ren, Jian & Xue, Yu. (2011). GPS- YNO2: Computational prediction of tyrosine nitration sites in proteins. Molecular bioSystems. 7. 1197–204. 10.1039/c0mb00279h.

2. Xu, Yan & Wen, Xin & Wen, Li-Shu & Wu, Ling-Yun & Deng, Nai-Yang & Chou, Kuo-Chen. (2014). iNitro-Tyr: Prediction of Nitrotyrosine Sites in Proteins with General Pseudo Amino Acid Composition. PloS one. 9. e105018. 10.1371/journal.pone.0105018.

3. Hasan, Md. Mehedi & Shamima, Khatun & Mollah, Md Nurul & Yong, Cao & Dianjing, Guo. (2018). NTyroSite: Computational Identification of Protein Nitrotyrosine Sites Using Sequence Evolutionary Features. Molecules. 23. 1667. 10.3390/molecules23071667.

4. Nilamyani, Andi & Auliah, Firda & Moni, Mohammad Ali & Shoombuatong, Watshara & Hasan, Md. Mehedi & Kurata, Hiroyuki. (2021). PredNTS: Improved and Robust Prediction of Nitrotyrosine Sites by Integrating Multiple Sequence Features. International Journal of Molecular Sciences. 22. 10.3390/ijms22052704.

5. Xie, Yubin & Luo, Xiaotong & Li, Yupeng & Chen, Li & Ma, Wenbin & Huang, Junjiu & Cui, Jun & Zhao, Yong & Xue, Yu & Zuo, Zhixiang & Ren, Jian. (2018). DeepNitro: Prediction of Protein Nitration and Nitrosylation Sites by Deep Learning. Genomics, Proteomics & Bioinformatics. 16. 10.1016/j.gpb.2018.04.007.

6. Naseer, Sheraz & Ali, Rao & Fati, Suliman & Muneer, Amgad. (2021). iNitroY-Deep: Computational Identification of Nitrotyrosine Sites to Supplement Carcinogenesis Studies Using Deep Learning. IEEE Access. PP. 1–1. 10.1109/ACCESS.2021.3080041.

7. Rahman, Afrida & Ahmed, Sabit & Hasan, Md. Al & Ahmad, Shamim & Dehzangi, Iman (Abdollah). (2022). Accurately Predicting Nitrosylated Tyrosine Sites Using Probabilistic Sequence Information. Gene. 826. 146445. 10.1016/j.gene.2022.146445.

8. Alberts, Bruce & Bray, Dennis & Hopkin, Karen & Johnson, Alexander D. & Lewis, Julian & Raff, Martin & Roberts, Keith & Walter, Peter. (2013). Essential Cell Biology. 4th ed. Philadelphia, PA: Garland Publishing.

9. Ramazi, Shahin & Zahiri, Javad. (2021). Posttranslational modifications in proteins: resources, tools and prediction methods. Database. 2021. 10.1093/database/baab012.

10. Xu, Haodong & Wang, Yongbo & Lin, Shaofeng & Deng, Wankun & Peng, Di & Cui, Qinghua & Xue, Yu. (2018). PTMD: A Database of Human Disease-associated Post-translational Modifications. Genomics, Proteomics & Bioinformatics. 16. 10.1016/j.gpb.2018.06.004.

11. Souza JM & Daikhin E & Yudkoff M & Raman CS & Ischiropoulos H. (1999). Factors determining the selectivity of protein tyrosine nitration. Archives of biochemistry and biophysics, 371(2), 169–78. 10.1006/abbi.1999.1480.

12. Abello, Nicolas & Kerstjens, Huib & Postma, Dirkje & Bischoff, Rainer. (2009). Protein Tyrosine Nitration: Selectivity, Physicochemical and Biological Consequences, Denitration, and Proteomics Methods for the Identification of Tyrosine-Nitrated Proteins. Journal of proteome research. 8. 3222–38. 10.1021/pr900039c.

13. Radi, Rafael. (2013). Protein tyrosine nitration: biochemical mechanisms and structural basis of functional effects. Accounts of chemical research, 46(2), 550–559. https://doi.org/10.1021/ar300234c

14. Li, Hailing & Yang, Zhen & Gao, Zhonghong. (2019). Protein tyrosine nitration: Chemistry and role in diseases. Advances in Molecular Toxicology. 10.1016/B978-0-444-64293-6.00004-X.

15. Chou, Kuo-Chen. (1993). A vectorized sequence-coupling model for predicting HIV protease cleavage sites in proteins. J. Biol. Chem. 268 (23), 16938–16948.

16. Chou, Kuo-Chen. (1995). A sequence-coupled vector-projection model for predicting the specificity of GalNAc-transferase. Protein Science 4: 1365–1383.

17. Chou, Kuo-Chen. (2001). Prediction of signal proteins using scaled window. proteins 22: 1973–1979.

18. Chou, Kuo-Chen. (2001). Prediction of protein cellular attributes using pseudo-amino acid composition. PROTEINS: Structure, Function, and Genetics (Erratum:ibid, 2001, Vol44, 60) 43: 246–255.

19. Ghandi, Mahmoud & Mohammad-Noori, Morteza & Beer, Michael. (2013). Robust k-mer frequency estimation using gapped k-mers. Journal of mathematical biology. 69. 10.1007/s00285-013-0705-3.

20. Ghandi, Mahmoud & Lee, Dongwon & Mohammad-Noori, Morteza & Beer, Michael. (2014). Enhanced Regulatory Sequence Prediction Using Gapped k-mer Features. PLoS computational biology. 10. e1003711. 10.1371/journal.pcbi.1003711.

21. Lyu, Zhibin & Zhang, Jun & Ding, Hui & Zou, Quan. (2020). RF-PseU: A Random Forest Predictor for RNA Pseudouridine Sites. Frontiers in Bioengineering and Biotechnology. 8. 10.3389/fbioe.2020.00134.

22. Manavalan, Balachandran & Hasan, Md. Mehedi & Basith, Shaherin & Gosu, Vijayakumar & Shin, Tae Hwan & Lee, Gwang. (2020). Empirical Comparison and Analysis of Web-Based DNA N 4 -Methylcytosine Site Prediction Tools. Molecular Therapy - Nucleic Acids. 22. 406–420. 10.1016/j.omtn.2020.09.010.

23. Almeida, Breno & Queiroz, Alvaro & Avila-Santos, Anderson & Bonidia, Robson & Nunes da Rocha, Ulisses & Sanches, Danilo & Carvalho, André. (2021). Feature Importance Analysis of Non-coding DNA/RNA Sequences Based on Machine Learning Approaches. 10.1007/978-3-030-91814-9_8.

24. Lyu, Zhibin & Wang, Donghua & Ding, Hui & Zhong, Bineng & Xu, Lei. (2020). Escherichia Coli DNA N-4-Methycytosine Site Prediction Accuracy Improved by Light Gradient Boosting Machine Feature Selection Technology. IEEE Access. PP. 1–1. 10.1109/ACCESS.2020.2966576.

25. Chen, Ke & Kurgan, Lukasz & Ruan, Jishou. (2007). Prediction of flexible/rigid regions from protein sequences using k-spaced amino acid pairs. BMC structural biology. 7. 25. 10.1186/1472-6807-7-25.

26. Zhao, Xiaowei & Zhang, Wenyi & Xu, Xin & Ma, Zhiqiang & Yin, Minghao. (2012). Prediction of Protein Phosphorylation Sites by Using the Composition of k-Spaced Amino Acid Pairs. PloS one. 7. e46302. 10.1371/journal.pone.0046302.

27. Hasan, Md. Mehedi & Zhou, Yuan & Lu, Xiaotian & Li, Jinyan & Song, Jiangning & Zhang, Ziding. (2015). Computational Identification of Protein Pupylation Sites by Using Profile-Based Composition of k-Spaced Amino Acid Pairs. PLoS ONE. 10. e0129635. 10.1371/journal.pone.0129635.

28. Hasan, Md. Mehedi & Kurata, Hiroyuki. (2018). GPSuc: Global Prediction of Generic and Species-specific Succinylation Sites by aggregating multiple sequence features. PLOS ONE. 13. e0200283. 10.1371/journal.pone.0200283.

29. Wei, Leyi & Tang, Jijun & Zou, Quan. (2017). SkipCPP-Pred: An improved and promising sequencebased predictor for predicting cell-penetrating proteins. BMC Genomics. 18. 1–11. 10.1186/s12864-017-4128-1.

30. Wei, Leyi & Xing, Pengwei & Su, Ran & Shi, Gaotao & Ma, Zhanshan & Zou, Quan. (2017). CPPred-RF: A Sequence-based Predictor for Identifying Cell-Penetrating proteins and Their Uptake Efficiency. Journal of Proteome Research. 16. 10.1021/acs.jproteome.7b00019.

31. Wei, Leyi & Xing, Pengwei & Shi, Gaotao & Ji, Zhi-Liang & Zou, Quan. (2017). Fast Prediction of Protein Methylation Sites Using a Sequence-Based Feature Selection Technique. IEEE/ACM Transactions on Computational Biology and Bioinformatics. PP. 1–1. 10.1109/TCBB.2017.2670558.

32. Wei, Leyi & Chen, Zhou & Chen, Huangrong & Song, Jiangning & Su, Ran. (2018). ACPred-FL: a sequencebased predictor using effective feature representation to improve the prediction of anti-cancer proteins. Bioinformatics. 34. 10.1093/bioinformatics/bty451.

33. Liu, Bin & Xu, Jinghao & Lan, Xun & Xu, Ruifeng & Zhou, Jiyun & Wang, Xiaolong & Chou, Kuo-Chen. (2014). iDNA-Prot—dis: Identifying DNA-Binding Proteins by Incorporating Amino Acid Distance-Pairs and Reduced Alphabet Profile into the General Pseudo Amino Acid Composition. PloS one. 9. e106691. 10.1371/journal.pone.0106691.

34. Liu, Bin & Gao, Xin & Zhang, Hanyu. (2019). BioSeq- Analysis2.0: an updated platform for analyzing DNA, RNA and protein sequences at sequence level and level based on machine learning approaches. Nucleic acids research. 47. 10.1093/nar/gkz740.

35. Ruiquan, Ge & Feng, Guanwen & Jing, Xiaoyang & Zhang, Renfeng & Wang, Pu & Wu, Qing. (2020). EnACP: An Ensemble Learning Model for Identification of Anticancer Peptides. Frontiers in Genetics. 11. 10.3389/fgene.2020.00760.

36. Li, Bo & Cai, Lijun & Liao, Bo & Fu, Xiangzheng & Bing, Pingping & Yang, Jialiang. (2019). Prediction of Protein Subcellular Localization Based on Fusion of Multi-view Features. Molecules. 24. 919. 10.3390/molecules24050919.

37. Wekesa, Jael & Meng, Jun & Luan, Yushi. (2020). Multi-feature fusion for deep learning to predict plant lncRNA-protein interaction. Genomics. 112. 2928–2936. 10.1016/j.ygeno.2020.05.005.

38. Singh, Dalwinder & Madhawan, Akansha & Roy, Joy. (2021). Identification of multiple RNAs using feature fusion. Briefings in Bioinformatics, Volume 22, Issue 6, bbab218, https://doi.org/10.1093/bib/bbab218

39. Jia, Yuran & Huang, Shan & Zhang, Tianjiao. (2021). KK- DBP: A Multi-Feature Fusion Method for DNA-Binding Protein Identification Based on Random Forest. Frontiers in Genetics. 12. 10.3389/fgene.2021.811158.

40. Guyon, Isabelle & Weston, Jason & Barnhill, Stephen & Vapnik, Vladimir. (2002). Gene Selection for Cancer Classification Using Support Vector Machines. Machine Learning. 46. 389–422. 10.1023/A:1012487302797.

41. Re, Matteo & Valentini, Giorgio. (2012). Ensemble methods: A review. Advances in Machine Learning and Data Mining for Astronomy, pp. 563–594. Chapman & Hall.

42. Schapire, Robert. (1990). The strength of weak learnability. Machine Learning. 5(2). 197–227. 10.1007/BF00116037

43. Wolpert, David. (1992). Stacked Generalization. Neural Networks. 5. 241–259. 10.1016/S0893-6080(05)80023-1.

44. Friedman, Jerome & Hastie, Trevor & Tibshirani, Robert. (2000). Additive Logistic Regression: A Statistical View of Boosting. The Annals of Statistics. 28. 337–407. 10.1214/aos/1016218223.

45. Friedman, Jerome. (2000). Greedy Function Approximation: A Gradient Boosting Machine. The Annals of Statistics. 29. 10.1214/aos/1013203451.

46. Friedman, Jerome. (2002). Stochastic Gradient Boosting. Computational Statistics & Data Analysis. 38. 367–378. 10.1016/S0167-9473(01)00065-2.

47. Cucchiara, Andrew. (2012). Applied Logistic Regression. Technometrics. 34. 358–359. 10.1080/00401706.1992.10485291.

48. Fawcett, Tom. (2006). Introduction to ROC analysis. Pattern Recognition Letters. 27. 861–874. 10.1016/j.patrec.2005.10.010.

49. Debie, Essam & Shafi, Kamran. (2019). Implications of the curse of dimensionality for supervised learning classifier systems: theoretical and empirical analyses. Pattern Analysis and Applications. 22. 10.1007/s10044-017-0649-0.

50. Weiss, Sophie & Xu, Zhenjiang & Peddada, Shyamal & Amir, Amnon & Bittinger, Kyle & González, Antonio & Lozupone, Catherine & Zaneveld, Jesse & Vázquez-Baeza, Yoshiki & Birmingham, Amanda & Hyde, Embriette & Knight, Rob. (2017). Normalization and microbial differential abundance strategies depend upon data characteristics. Microbiome. 5. 10.1186/s40168-017-0237-y.

51. Abrams, Zachary & Johnson, Travis & Huang, Kun & Payne, Philip & Coombes, Kevin. (2019). A protocol to evaluate RNA sequencing normalization methods. BMC Bioinformatics. 20. 10.1186/s12859-019-3247-x.

52. Lovell, David & Müller, Warren & Taylor, Jennifer & Zwart, Alec & Helliwell, Chris. (2011). Proportions, Percentages, PPM: Do The Molecular Biosciences Treat Compositional Data Right?. 10.1002/9781119976462.ch14.

53. Breiman, Leo. (2000). Bias, Variance, And Arcing Classifiers. Technical Report 460, Statistics Department, University of California.

54. Edgar, Robert. (2010). Search and Clustering Orders of Magnitude Faster than BLAST. Bioinformatics (Oxford, England). 26. 2460–1. 10.1093/bioinformatics/btq461.

55. Fu, Limin & Zhu, Zhengwei & Wu, Sitao & Li, Weizhong. (2012). CD-HIT: Accelerated for clustering the next-generation sequencing data. Bioinformatics (Oxford, England). 28. 10.1093/bioinformatics/bts565.

56. Natekin, Alexey & Knoll, Alois. (2013). Gradient Boosting Machines, A Tutorial. Frontiers in neurorobotics. 7. 21. 10.3389/fnbot.2013.00021.

57. Mishra, Sidharth & Sarkar, Uttam & Taraphder, Subhash & Datta, Sanjoy & Swain, Devi & Saikhom, Reshma & Panda, Sasmita & Laishram, Menalsh. (2017). Principal Component Analysis. International Journal of Livestock Research. 1. 10.5455/ijlr.20170415115235.

58. van der Maaten, Laurens & Hinton, Geoffrey. (2008). Viualizing data using t-SNE. Journal of Machine Learning Research. 9. 2579–2605.

59. Demir-Kavuk, Ozgur & Kamada, Mayumi & Akutsu, Tatsuya & Knapp, Ernst-Walter. (2011). Prediction using step-wise L1, L2 regularization and feature selection for small data sets with large number of features. BMC bioinformatics. 12. 412. 10.1186/1471-2105-12-412.

60. Tibshirani, Robert. “Regression Shrinkage and Selection via the Lasso.” Journal of the Royal Statistical Society. Series B (Methodological), vol. 58, no. 1, 1996, pp. 267–88. JSTOR, http://www.jstor.org/stable/2346178. Accessed 29 Nov. 2022.

61. Hoerl, Arthur E., & Robert W. Kennard. “Ridge Regression: Biased Estimation for Nonorthogonal Problems.” Technometrics, vol. 42, no. 1, 2000, pp. 80–86. JSTOR, https://doi.org/10.2307/1271436. Accessed 29 Nov. 2022.

62. Hornbeck, Peter & Kornhauser, Jon & Tkachev, Sasha & Zhang, Bin & Skrzypek, Elzbieta & Murray, Beth & Latham, Vaughan & Sullivan, Michael. (2011). PhosphoSitePlus: A comprehensive resource for investigating the structure and function of experimentally determined post-translational modifications in man and mouse. Nucleic acids research. 40. D261–70. 10.1093/nar/gkr1122.

63. Matlock, Matthew & Holehouse, Alex & Naegle, Kristen. (2014). ProteomeScout: A repository and analysis resource for post-translational modifications and proteins. Nucleic acids research. 43. 10.1093/nar/gku1154.

64. Peri, S. et al. (2003). Development of Human Protein Reference Database as an initial platform for approaching systems biology in humans. Genome Research. 13, 2363–2371.

65. Sigrist, Christian & Cerutti, Lorenzo & Castro, Edouard & Langendijk-Genevaux, Petra & Bulliard, Virginie & Bairoch, Amos & Hulo, Nicolas. (2009). PROSITE, a protein domain database for functional characterization and annotation. Nucleic Acids Res 38:D161-D166. Nucleic acids research. 38. D161–6. 10.1093/nar/gkp885.

66. Wu, Cathy & Yeh, Lai-Su & Huang, Hongzhan & Arminski, Leslie & Castro-Alvear, Jorge & Chen, Yongxing & Hu, Zhangzhi & Kourtesis, Panagiotis & Ledley, Robert & Suzek, Baris & Vinayaka, C. R. & Zhang, Jian & Barker, Winona. (2003). The Protein Information Resource. Nucleic acids research. 31. 345–7. 10.1093/nar/gkg040.

67. Li, Zhongyan & Li, Shangfu & Luo, Mengqi & Jhong, Jhih-Hua & Li, Wenshuo & Yao, Lantian & Pang, Yuxuan & Wang, Zhuo & Wang, Rulan & Ma, Renfei & Yu, Jinhan & Huang, Yuqi & Zhu, Xiaoning & Cheng, Qifan & Feng, Hexiang & Zhang, Jiahong & Wang, Chunxuan & Hsu, Justin & Chang, Wen-Chi & Lee, Tzong-Yi. (2021). dbPTM in 2022: an updated database for exploring regulatory networks and functional associations of protein post-translational modifications. Nucleic acids research. 50. 10.1093/nar/gkab1017.

68. Li, Jing & Jia, Jia & Li, Hong & Yu, Jian & Sun, Han & He, Ying & Lv, Daqing & Yang, Xiaojuan & Glocker, Michael & Ma, Liangxiao & Yang, Jiabei & Li, Ling & Li, Wei & Zhang, Guoqing & Liu, Qian & Li, Yixue & Xie, Lu. (2014). SysPTM 2.0: An updated systematic resource for post-translational modification. Database: the journal of biological databases and curation. 2014. bau025. 10.1093/database/bau025.

69. Bateman, Alex & Martin, Maria-Jesus & Orchard, Sandra & Magrane, Michele & Agivetova, Rahat & Ahmad, Shadab & Alpi, Emanuele & Bowler-Barnett, Emily & Britto, Ramona & Bursteinas, Borisas & Bye-A-Jee, Hema & Coetzee, Ray & Cukura, Austra & Silva, Alan & Denny, Paul & Dogan, Tunca & Ebenezer, Thankgod & Fan, Jun & Castro, Leyla & Teodoro, Douglas. (2020). UniProt: the universal protein knowledgebase in 2021. Nucleic Acids Research. 49. 10.1093/nar/gkaa1100.

70. Wulff-Fuentes, Eugenia & Berendt, Rex & Massman, Logan & Danner, Laura & Malard, Florian & Vora, Jeet & Kahsay, Robel & Olivier-Van Stichelen, Stéphanie. (2021). The human O-GlcNAcome database and meta-analysis. Scientific Data. 8. 10.1038/s41597-021-00810-4.

71. Liashchynskyi, Petro & Liashchynskyi, Pavlo. (2019). Grid Search, Random Search, Genetic Algorithm: A Big Comparison for NAS. 10.48550/arxiv.1912.06059.

72. Bonidia, Robson & Domingues, Douglas & Sanches, Danilo & Carvalho, André. (2021). MathFeature: feature extraction package for DNA, RNA and protein sequences based on mathematical descriptors. Briefings in Bioinformatics. 23. 10.1093/bib/bbab434.

73. Chen, Zhen & Zhao, Pei & Li, Chen & Li, Fuyi & Xiang, Dongxu & Chen, yongzi & Akutsu, Tatsuya & Daly, Roger & Webb, Geoffrey & Zhao, Quanzhi & Kurgan, Lukasz & Song, Jiangning. (2021). iLearnPlus: A comprehensive and automated machine-learning platform for nucleic acid and protein sequence analysis, prediction and visualization. Nucleic Acids Research. 49. e60. 10.1093/nar/gkab122.

74. Chen, Tianqi & Guestrin, Carlos. (2016). XGBoost: A Scalable Tree Boosting System. 785–794. 10.1145/2939672.2939785.

75. Pedregosa, Fabian & Varoquaux, Gael & Gramfort, Alexandre & Michel, Vincent & Thirion, Bertrand & Grisel, Olivier & Blondel, Mathieu & Prettenhofer, Peter & Weiss, Ron & Dubourg, Vincent & Vanderplas, Jake & Passos, Alexandre & Cournapeau, David & Brucher, Matthieu & Perrot, Matthieu & Duchesnay, Edouard & Louppe, Gilles. (2012). Scikit-learn: Machine Learning in Python. Journal of Machine Learning Research. 12, 2825–2830.

76. Tareen, Ammar & Kinney, Justin. (2019). Logomaker: Beautiful sequence logos in Python. Bioinformatics (Oxford, England). 36. 10.1093/bioinformatics/btz921.

